# Multimodal analysis unveils tumor microenvironment heterogeneity linked to immune activity and evasion

**DOI:** 10.1101/2023.12.20.572033

**Authors:** Óscar Lapuente-Santana, Gregor Sturm, Joan Kant, Markus Ausserhofer, Constantin Zackl, Maria Zopoglou, Nicholas McGranahan, Dietmar Rieder, Zlatko Trajanoski, Noel Filipe da Cunha Carvalho de Miranda, Federica Eduati, Francesca Finotello

## Abstract

The cellular and molecular heterogeneity of tumors is a major obstacle to cancer immunotherapy. Here, we use a systems biology approach to derive a signature of the main sources of heterogeneity in the tumor microenvironment (TME) from lung cancer transcriptomic data. We demonstrate that this signature, which we called *iHet*, is conserved in different cancers and associated with antitumor immunity. Through the analysis of single-cell and spatial transcriptomics data, we trace back the cellular origin of the variability that explains the iHet signature. Finally, we demonstrate that iHet has predictive value for cancer immunotherapy, which can be further improved by disentangling three major determinants of anticancer immune responses: activity of immune cells, immune infiltration or exclusion, and cancer-cell foreignness. This work shows how transcriptomics data can be integrated to derive a holistic representation of the phenotypic heterogeneity of the TME, and ultimately to determine its unfolding and fate during immunotherapy with immune checkpoint blockers.

## Introduction

In recent years, immunotherapies have emerged as powerful anticancer treatments, transforming the practice of medical oncology and bringing new hope to patients with different cancers^1^. Instead of directly targeting tumor cells, immunotherapy harnesses the body’s own immune system to fight cancer. Immune checkpoint blockers (ICB) have been the major drivers of cancer immunotherapy revolution and, to date, represent the most successful treatment regimen for advanced cancers^2,3^. Immune checkpoint blockade relies on monoclonal antibodies that target immune-cell regulators like cytotoxic T-lymphocyte-associated antigen-4 (CTLA-4) and programmed cell death protein 1 (PD-1) to elicit or boost antitumor immunity. ICB have shown unprecedented long-term clinical efficacy and are now part of the standard of care for patients with different cancer types^2–4^.

Nevertheless, patients’ responses to ICB remain limited and hard to predict. The main obstacles stem from our incomplete understanding of how anticancer immune responses are orchestrated or regulated within the tumor microenvironment (TME)^5^. This complexity is exacerbated by the inherent cellular and molecular heterogeneity of the TME, which complicates the development of effective anticancer treatments^6,7^. Clinical studies have revealed three major cancer-immune phenotypes associated with patients’ responses to ICB immunotherapy (reviewed in ^8–11)^. The *immune-inflamed* (or *immune*-*enriched*) phenotype is characterized by the presence of large numbers of effector T cells in close contact with cancer cells. Clinical responses to ICB occur more often in patients with immune-inflamed tumors (also called “hot” tumors) but are not warranted, indicating that additional factors besides immune-cell infiltration play a role in inducing a therapeutic response^8^. In the *immune-excluded* phenotype, immune cells are kept at the border of the tumor, restricted to the tumor stroma. The likelihood of clinical response to ICB is more limited in these tumors as compared to the immune inflamed ones^8^. Finally, *immune-desert* tumors are characterized by limited or absent T-cell infiltration. Myeloid cells can be present, but their general feature is a non-inflamed TME (“cold” tumors) and infrequent responses to immunotherapy^8^.

However, this broad classification does not fully recapitulate more nuanced differences between tumors (i.e. inter-tumor heterogeneity) nor the geospatial heterogeneity of tumors (i.e. intra-tumor heterogeneity), failing to explain the interwoven molecular and cellular mechanisms that ultimately determine ICB response. When speaking about intra-tumor heterogeneity, the genetic diversity of malignant cells is probably its best understood component^12,13^. Nevertheless, the *phenotypic* diversity of tumors, which is the object of this study, is the result of a much more complex and dynamic entanglement of the epigenetic, transcriptomic, metabolic, and communication programmes of cancer, immune, stromal, and endothelial cells that compose the TME^6,7^.

The analysis of tumor transcriptomic data has strong potential for investigating the facets of tumor heterogeneity influencing patients’ response to immunotherapy. However, distilling large-scale information from RNA-seq studies, from thousands of measured genes towards a few interpretable features that are valuable for clinical decision support, is far from trivial. One common approach is to focus on processes known to be associated with anticancer immunity (e.g. cytolytic activity of tumor-infiltrating lymphocytes^14^, presence of tertiary lymphoid structures^15^, IFN-γ signaling^16^) and derive a related gene signature that can be used to predict patients’ response to immunotherapy from pre-therapy RNA-seq data. Despite their predictive value for immunotherapy, these gene-centric approaches have intrinsic limitations that narrow their potential to enhance our understanding of how anticancer immune responses are orchestrated. First, gene signatures have limited interpretability and hardly provide new mechanistic insights on the actual molecular and cellular processes underlying immunological control. Second, they usually adopt a univariate approach, modeling single processes selected based on prior knowledge and, thus, fail to provide a holistic view on how pro- and anti-tumor mechanisms dynamically integrate in the TME. Third, as they are based solely on bulk RNA-seq data analysis, they might fail in capturing how intra-tumor heterogeneity influences the success of cancer immunotherapy.

In this study, we took a systems biology approach, using tumor RNA-seq data to extract an interpretable, high-level (rather than gene-centric) representation of the TME in terms of cell types, pathways, and transcription factors. For this purpose, we used publicly-available, bulk transcriptomic data, generated from non-small cell lung cancer tumors (1094 patients and 1244 samples). We chose to focus on non-small cell lung cancer (NSCLC) for its marked intra-tumor heterogeneity, encompassing not only the genetic makeup of cancer cells but also the overall cellular composition of the TME and its immune contexture^17–19^. Moreover, NSCLC tumor RNA-seq data is available from large cohorts of patients as well as from multi-biopsy transcriptomic studies, allowing the investigation of both intra- and inter-tumor heterogeneity. Starting from this high-level, multimodal data, we used an unsupervised approach to provide a low-dimensional representation of the data and identify the latent factor capturing most of the data variance. We showed, *a posteriori*, that this factor is associated with immune response. This latent factor, that we called *immune heterogeneity (iHet)* signature, gives a quantitative, interpretable, and holistic representation of TME cells, pathways, and transcription factors that are associated with immune infiltration and activity, but also with negative feedback mechanisms underlying immune evasion. Through the extension of our analysis to additional 16 cancer types, with more than 6000 bulk-tumor RNA-seq samples analyzed, we demonstrate how this holistic signature is conserved in different solid cancers and can be used to predict patients’ response to ICB immunotherapy. Finally, we showed how the integration of genomic and digital pathology data provides orthogonal information on facets of the TME that are not captured by iHet, disentangling the determinants of anticancer immunity from mechanisms of immune evasion. Notably, the combination of three complementary axes quantifying immune response, immune exclusion, and tumor foreignness augmented the predictive value of iHet for ICB immunotherapy in different solid cancers.

## Results

### Unsupervised analysis identifies a conserved signature associated with phenotypic heterogeneity and immune response in NSCLC

To investigate the phenotypic heterogeneity of NSCLC tumors, we analyzed RNA-seq data generated from multi-region biopsies collected in two recent studies (referred to as ‘Jia’^18^ and ‘Sharma’^19^ hereafter), as well as from bulk tumors of lung adenocarcinoma (LUAD) and lung squamous cell carcinoma (LUSC) patients from The Cancer Genome Atlas (TCGA)^20^; dataset details in **Supplementary Table 1**. In order to gain a high-level view of the sources of heterogeneity in NSCLC, we extracted from RNA-seq data quantitative information on the cellular composition of the tumor microenvironment (TME) via deconvolution with quanTIseq^21^ and EPIC^22^, as well as on pathway and transcription factor (TF) activity using PROGENy^23^ and DoRothEA^24^, respectively (**STAR Methods**). Deconvolution methods leverage cell-type-specific signatures to infer the cellular composition of bulk RNA-seq samples^25^, whereas “footprint” methods like PROGENy and DoRothEA can infer from the same data the activity of pathways and TF by considering the expression of genes regulated by the process of interest (e.g., pathway-responsive genes or TF targets, respectively)^26^. The final data encompassed 10 cell types, 14 pathways, and 118 TFs, which can be seen as multimodal features.

After normalization and integration of these features, we performed unsupervised Multi-Omics Factor Analysis (MOFA)^27^ to disentangle the major axes of heterogeneity (**Figure 1A** and **STAR Methods**). MOFA can be seen as a generalization of principal component analysis (PCA) for multimodal data. MOFA seeks to identify latent factors (F1, F2, F3, …), computed as linear combinations of the input features, which capture the major sources of variation across data modalities (**Figure 1A**). We applied MOFA to the Jia and Sharma multi-biopsy datasets, as well as to the LUAD and LUSC TCGA datasets, analyzed both singularly and jointly (‘JiaSharma’ and ‘TCGA-NSCLC’). MOFA was run in bootstrap settings to robustly identify latent factors capturing the major sources of intra- and inter-patient variance (**STAR Methods**). We observed that the resulting latent factor F1, which explained most of the data variance across the three modalities (**Supplementary Figure 1**), was conserved in the different NSCLC datasets (**Figure 1B**, Pearson correlations of the F1 weights for the single and joint datasets ranging in r = 0.75-0.98). As a control, we ran MOFA also on RNA-seq data generated from healthy lung tissues by the Genotype-Tissue Expression (GTEx) initiative^28^. We observed only a weak correlation between the F1 weights extracted from the GTEx and NSCLC datasets (r = 0.0-0.52) (**Figure 1C** and **Supplementary Figure 2**), indicating its tumor specificity.

**Figure 1.**
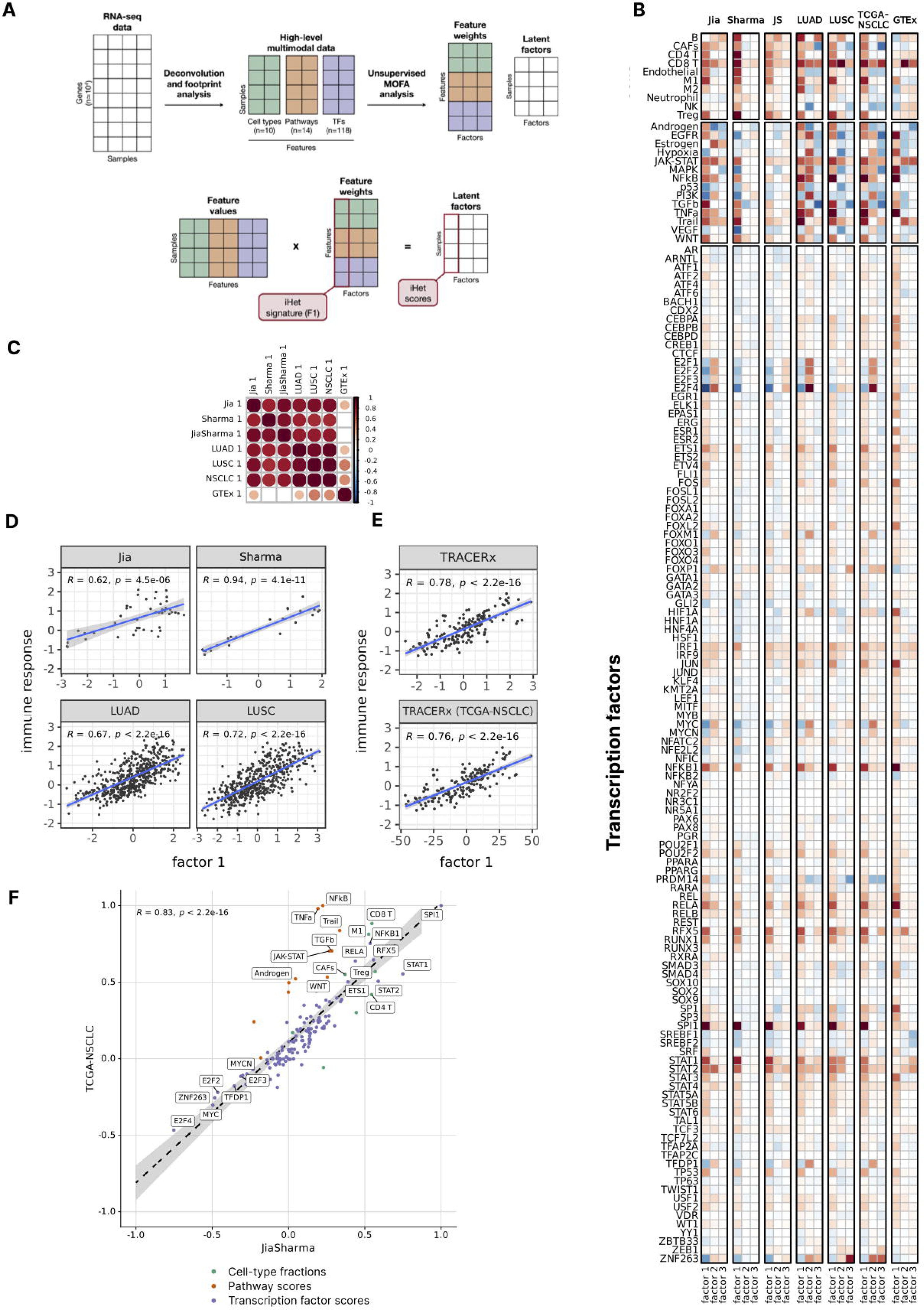
Identification of an immune heterogeneity (iHet) signature via unsupervised multimodal analysis. **(A)** Schematization of the analytical methodology. Large-scale RNA sequencing (RNA-seq) data is processed with deconvolution and footprint methods to estimate cell type abundances, together with pathway and transcription factor (TF) activities. This “high-level” multimodal data is jointly analyzed using Multi-Omics Factor Analysis (MOFA) to derive latent factors (F1, F2, …) capturing most of the data variance and the factor weights associated to each feature. The F1 latent factor represents the *immune heterogeneity* (iHet) score across all patients and can be calculated via matrix multiplication of the feature values with the F1 feature weights (iHet signature). **(B)** Heatmap of the median weights for MOFA F1-F3 latent factors across 100 bootstrap runs for the lung-cancer and healthy-lung datasets, separated by feature modality. **(C)** Pearson correlation of F1 weights across datasets. Dot sizes correspond to the absolute correlation coefficients. Only dots with false-discovery rate (FDR)-adjusted p-values < 0.01 are shown. **(D)** Correlation of the F1 factor with the ensemble immune response score, derived from state-of-the-art prediction methods, for LUAD, LUSC, Jia, Sharma datasets. **(E)** Correlation of TRACERx F1 factor (estimated by MOFA) and iHet (estimated by multiplying the features scores from the TRACERx dataset with the F1 weights derived from the TCGA-NSCLC dataset) with the ensemble immune response score. **(F)** Scatter plot of F1 weights derived from the TCGA-NSCLC and JiaSharma combined datasets, respectively, coloured by feature type. Top features with weights > 0.5 or < -0.3 on either of the two axes are labeled. In all scatterplots, *R* represents Pearson correlation, *p* the associated, two-tailed p-value.

As this conserved axis of heterogeneity identified with our unsupervised approach captured several immune cells involved in anticancer immunity (**Figure 1B**), we decided to assess its association with immune response. To derive an immune response score, we used EaSIeR^29^ to integrate nine state-of-the-art methods that have been used to predict patients’ response to immunotherapy with ICB from pre-therapy RNA-seq data^14–16,30–33^. Briefly, we scaled the nine scores to a comparable range of values through z-scoring, and derived an ensemble immune response score as the median of their z-scores (details in **STAR Methods**). Correlation analysis revealed that feature weights associated with the conserved latent factor F1, despite being identified with an unsupervised analysis, were strongly positively associated with the predicted immune response in all NSCLC datasets (**Figure 1D**). To validate these findings in an independent cohort, we considered an RNA-seq dataset from 164 multi-region biopsies collected from 64 NSCLC patients of the TRACERx study^17^. We derived the F1 latent factor for each sample in two alternative ways. First, we performed unsupervised MOFA analysis and extracted the factor explaining most of the data variance from the TRACERx dataset, as done for the other NSCLC datasets. In parallel, we computed the TRACERx latent factor F1 using the feature weights previously derived for TCGA-NSCLC (**STAR Methods; Figure 1A**). Both types of F1 were also strongly, positively associated with the predicted immune response in the TRACERx cohort (**Figure 1E**).

Taken together, these results suggest that our unsupervised multimodal analysis identified a conserved and interpretable signature underlying tumor phenotypic heterogeneity, which is associated with (predicted) immune response in multiple NSCLC cohorts. Hereafter, we will refer to the F1 feature weights derived via MOFA analysis from the TCGA-NSCLC data as *immune heterogeneity (iHet) signature* (**Supplementary Table 2**), and to the per-sample latent factor derived via matrix multiplication of these weights by the sample-specific feature values as *iHet score*.

### iHet top features describe the cellular and molecular underpinnings of NSCLC phenotypic heterogeneity

We compared the median F1 feature weights for the joint MOFA models derived from the JiaSharma multi-biopsy and TCGA-NSCLC datasets (**Figure 1F**). Despite being derived in an unsupervised manner from different datasets covering different types of heterogeneity, the two sets of iHet feature weights were in strong agreement (Pearson correlation of 0.83, p < 2.2·10^-16^). The feature with the largest positive weight –and, thus, positive association with the immune response score– in both JiaSharma and TCGA-NSCLC models was SPI1. SPI1 encodes the transcription factor PU.1, which is a major transcriptional regulator within the hematopoietic system, required for myeloid and B-lymphoid cell development^34^. The E2F4 transcription factor had the largest negative weight in both models. E2F4 is part of the E2F family of transcription factors, which play a key role in the control of cell cycle and act as tumor suppressor proteins^35^. Among the features with positive weight in both datasets, there were immune cells associated with a good clinical outcome like CD8^+^ T cells and M1 macrophages^36,37^, as well as markers of interferon gamma (IFN-γ) response, such as JAK-STAT pathway as well as STAT1 and STAT2 transcription factors^38^. Other positive weights were identified for molecular and cellular components associated with immune-suppressive mechanisms, like regulatory T (T_REG_) cells, cancer-associated fibroblasts (CAF), and TGFβ pathway.

The weights of the top features obtained from the different NSCLC models (labeled in **Figure 1F**) showed limited variability across bootstrap runs, especially for TF, and systematically positive or negative values with the only exception of the Androgen pathway (**Supplementary Figure 4A**). Overall, these further support the robustness of the iHet signature. The within-dataset variability was higher in the multi-biopsy studies compared to TCGA data, likely due to their smaller sample size and to the multiregional nature of the data. Leveraging the three multi-biopsy RNA-seq datasets, we could also systematically assess the intra- and inter-patient variability of iHet features. To this end, we computed a stability score to quantify, for each feature, how stable were patients’ ranks across multiple biopsies. The stability score was derived by computing multiple times the rankings of patients for each feature, quantified each time using a single patient’s biopsy selected through random sampling (**STAR Methods**). Features with high stability scores were E2F2, MYC, and MYCN transcription factors, while JAK-STAT was the most stable pathway (**Supplementary Figure 4B**). On the contrary, CAF abundance together with the activity of Trail, TGFβ, and TNFα pathways were among the most variable intra-patient. Of note, the latter features were also the ones showing lower weights in the JiaSharma model compared to the TCGA-NSCLC one (**Figure 1F**), which could stem from their higher variability in the multi-biopsy data. Overall, the results were dependent on the specific dataset. In particular, the stability of CD8^+^ T cell abundance was variable across datasets, likely due to differences in immune infiltration in the different tumors. We confirmed these results using an alternative approach: the ratio between inter- and intra-patient heterogeneity (details in **STAR Methods**) showed a positive Pearson correlation with the stability score (0.64 for Jia, 0.72 for Sharma, and 0.66 for TRACERx, p < 2.2·10^-16^). Notably, intra-sample transcriptional variability had also an impact on iHet scores computed for each biopsy, which was extensive in some tumors (e.g. Jia-P022) and more limited in others (e.g. Jia-P009) (**Supplementary Figure 5**).

### Analysis of single-cell and spatial transcriptomic data reveals the cellular origin of iHet

To investigate the cellular sources of the signal captured by the iHet signature (i.e. the weights of the F1 factor identified through unsupervised analysis of TCGA-NSCLC data), we reconstructed the TME of NSCLC at single-cell resolution leveraging our previously published lung cancer atlas (LuCA)^39^. From this compendium of data, we selected only tumor samples (i.e., no healthy tissue) annotated as either LUAD or LUSC, finally integrating 801,488 single-cell RNA-seq (scRNA-seq) profiles from 208 NSCLC patients across 21 datasets^39–49^ (**Supplementary Table 1** and **Supplementary Figure 6**). Briefly, after quality control, doublet detection, and harmonization of gene symbols and metadata, we trained a deep generative model using scANVI^50^ to perform data integration (**STAR Methods**). We carried out unsupervised, graph-based clustering and used the expression of cell type-specific marker-genes to manually assign cells to twelve major cell populations (**Figure 2A** and **Supplementary Figure 6A**).

**Figure 2.**
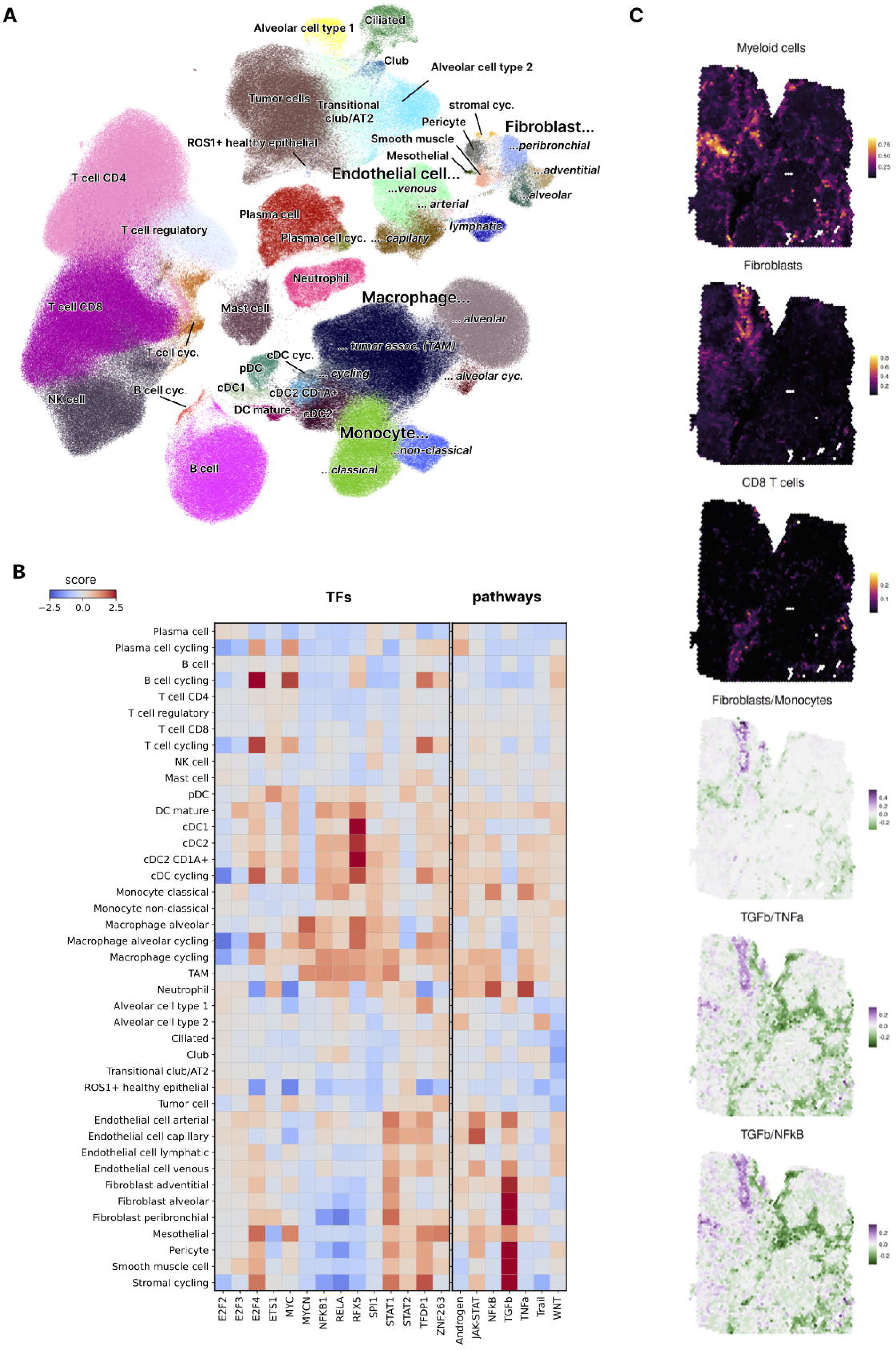
Cellular origin of of iHet unveiled by single-cell and spatial transcriptomics data analysis. **(A)** UMAP plot of the full NSCLC single-cell atlas, coloured by cell-type. **(B)** Mean activity scores per cell type for the top transcription factors (TF, left panel) and pathways (left panel), as labeled in Figure 1F. Scores are clipped at -2.5 and 2.5, respectively. **(C)** Exemplary spatial transcriptomics analysis of a lung cancer slide profiled with the 10x Visium technology. The first three panels show the estimated cell-type fractions per spot for myeloid cells, fibroblasts, and CD8^+^ T cells, respectively. The last three panels display the log-ratio of the estimated peribronchial fibroblast vs. classical monocyte cell fractions, the TGFβ vs. TNFα pathway scores, and the TGFβ vs. NF-κB pathway scores, respectively.

We iteratively subdivided these populations using graph-based clustering at a higher resolution and annotated more fine-grained cell subsets based on known marker genes (**Supplementary Figure 7**). Among stromal cells, T cells, dendritic cells (DC), plasma cells, and macrophages, subsets of cycling cells could be identified thanks to the expression of cell-cycle and proliferation markers like *CDK1* and *MKI67*. We distinguished malignant epithelial cells from healthy epithelial cells, and disentangled four types of endothelial cells: venous, arterial, capillary, and lymphatic. The stromal compartment was mainly composed of fibroblasts (peribronchial, adventitial, and alveolar), together with pericytes, smooth muscle cells, and mesothelial cells. Among the lymphocytic compartment, we could identify CD8^+^ T cells, effector CD4^+^ T cells, regulatory T_REG_ cells, and natural killer (NK) cells, plus a well-separated cluster of B cells. Also neutrophils, mast cells, plasma cells, and IL3RA-expressing plasmacytoid DC (pDC) constituted each a separated cluster.

The myeloid compartment constituted the largest cluster (198,237 cells, 25% of total cells) after the T/NK-compartment (337,205 cells, 42% of total cells) and it was characterized by a marked diversity in terms of DC, monocyte, and macrophage subtypes (**Supplementary Figure 8A-B**). We identified a cluster of classical monocytes characterized by the expression of *CD14* and *VCAN*, and a cluster of non-classical monocytes expressing *FCGR3A* and *LST1*. Myeloid DC could be divided into five subclusters, including a subset of cycling DC. Mature DC, which showed a clear separation from the other classical DC (cDC) subsets (**Supplementary Figure 8A-B**), expressed the maturation markers *CCR7*, *CD40*, *RELB*, *CD83*, together with *LAMP3* and, to a lesser extent, *CD274* (*PDL1*). cDC1 also segregated in a separate cluster, characterized by the expression of *CLEC9A*, *XCR1*, and *IRF8*. Finally, besides classical cDC2 expressing *CD1C* and *FCER1A*, we could identify a subpopulation of “langerhans-like” DC expressing also *CD207*, which encodes the Langerin transmembrane protein, and *CD1A*. All myeloid subtypes were represented by scRNA-seq data from multiple patients and datasets (**Supplementary Figure 8C).**

To disentangle the cellular contribution to the top molecular features associated to the iHet signature, we quantified pathway and TF activity in single cells using the same computational tools used for the bulk RNA-seq data, namely PROGENy and Dorothea, selected for their good performance also on scRNA-seq data^51^. Most of the pathways and TFs of interest showed activity in more than one cell type (**Figure 2B** and **Supplementary Table 3**). Many pathways and TFs with positive iHet weights showed high activity in myeloid cells, including SPI1, TNFα, MYCN, NF-κB, and RFX5 (**Figure 2B-C** and **Supplementary Fig 9**). In particular, SPI1 showed high activity in all myeloid subsets except cDC1 dendritic cells. NF-κB and TNFα also showed extensive activity across myeloid subsets, with the exception of cDC1, mature DC, and non-classical monocytes. Other features showed instead an enrichment in specific myeloid subsets, like MYCN (TAM and alveolar macrophages) and RFX5 (myeloid DC, especially cDC1, cDC2, and “langerhans-like” cD1A^+^ cDC2). TGFβ showest the highest activity in fibroblasts and other stromal cell subsets. As expected, due to their role in the regulation of gene transcription during cell cycle progression, E2F4 and TFDP1 showed higher activity in cycling cells across different lineages, but also higher activity in a subset of malignant cells (**Figure 2C**).

Taken together, our analysis of bulk and single-cell RNA-seq data indicated a major contribution of myeloid cells and fibroblasts to tumor and immune heterogeneity, as well as regional variability of these two cellular compartments within the same TME. Despite the limited availability of spatially-resolved data from the NSCLC TME, we decided to show an example of how these results can be further validated by analyzing a publicly-available spatial transcriptomics dataset generated from a lung cancer slide using the 10x Genomics Visium platform (www.10xgenomics.com). We used the spacedeconv toolkit (manuscript in preparation) to determine the spatial organization and most active pathways of myeloid cells and fibroblasts (details in **STAR Methods**). Briefly, after data preprocessing and normalization, we quantified the cellular composition of each spot using cell2location^52^ trained with Lambrechts scRNA-seq data^42^ (which constituted 5% of our single-cell atlas). decoupleR^53^ was instead used to quantify TNFα and NF-κB pathways, which according to our single-cell analysis were most active in myeloid cells –especially in classical monocytes– and TGFβ pathway, most active in fibroblasts (**Figure 2C** and **Supplementary Figure 6E**). Myeloid cells and fibroblasts segregated in specific regions of the tumor rather than being distributed across the whole slide. Among myeloid cells, classical monocytes constituted the most abundant subset (average cell fractions across all spots equal to 0.05), followed by tumor-associated macrophages (TAMs, 0.03). Among fibroblasts, peribronchial fibroblasts were the most abundant subset (0.04), followed by adventitial fibroblasts (<0.01). We assessed in more detail the relative locations of classical monocytes and peribronchial fibroblasts in the tumor using spacedeconv (**Figure 2C**). Fibroblasts and classical monocytes were closely colocalized in two regions located in the top-left corner of the slide. The localization of monocyte and fibroblasts was further supported by the activity of monocyte-specific (TNFα and NF-κB) and fibroblast-specific (TGFβ) pathways, corroborating the pathway-cell type associations identified through our single-cell analysis.

Taken together, these results show how scRNA-seq and spatial transcriptomics data can be used to identify the cellular origins of TME heterogeneity, together with their underlying molecular programs. In line with our bulk and multi-biopsy RNA-seq data analyses, our analysis of single-cell and spatial transcriptomics indicates a high abundance of the myeloid and fibrotic components, and suggests their spatial variability within the same TME.

### Disentangling immune activity and exclusion improves the prediction of ICB response in different cancer types

The results of our bulk and single-cell analyses of NSCLC transcriptomics show that the iHet signature, identified with an unsupervised computational approach, encompasses molecular and cellular underpinnings of anticancer immune responses. This suggests that the iHet score could be used to predict the likelihood of response of patients’ to ICB immunotherapy. However, our results also show that iHet captures mechanisms associated with immune suppression and, eventually, with immune exclusion, like CAF abundance and TGFβ pathway activity. In this light, it is key to disentangle these two opposing iHet components, in order to distinguish immune-inflamed tumors from immune-excluded ones, where the latter are less likely to respond to ICB^8,54^.

To systematically test which iHet features are associated with immune exclusion, we leveraged orthogonal information extracted from digital pathology tumor data. Since fibrosis is one of the key factors that leads to immune exclusion^55^, we considered fibroblast density estimates inferred from hematoxylin and eosin (H&E)-stained slides of TCGA NSCLC patients^56^ (**STAR Methods**). Through correlation analysis, we verified that 64 out of the 142 iHet features were positively correlated (Spearman correlation r > 0, FDR ≤ 0.05) with the density of fibroblasts in the tumor region (**Figure 3A** and **Supplementary Table 4**). The top five features associated with fibroblast density were JUN (enriched in T-cell single-cell data, **Supplementary Table 3**), SMAD4 and SMAD3 transcription factors, CAFs, and TGFβ pathway. The identification of molecularly-defined features reflecting CAF presence, together with their associated activity and regulatory circuitries, such as TGFβ pathway and SMAD3/SMAD4 transcription factors^57^ (also enriched in CAFs single-cell data, **Supplementary Table 3**), endorses the validity of our data integration approach. To further verify the association of these 64 features with fibroblast-mediated immune exclusion, we evaluated these results in the context of the cancer-immune phenotypes defined in a recent study^58^, where the authors used TCGA RNA-seq data across different TCGA tumor types, including LUAD and LUSC, to derive four TME classes: desert (“D”), fibrotic (“F”), immune enriched (“IE”), and immune enriched fibrotic (“IE/F”)^58^. Through this analysis, we revealed that 52/64 (i.e. 81%) of the features that we defined as associated with immune exclusion based on their correlation with imaging-derived fibroblast density also had higher values in immune-enriched fibrotic (“IE/F”) or fibrotic (“F”) tumors compared to the immune-enriched (“IE”) ones (**Figure 3A** and **Supplementary Figure 10**, details in **STAR Methods**).

**Figure 3.**
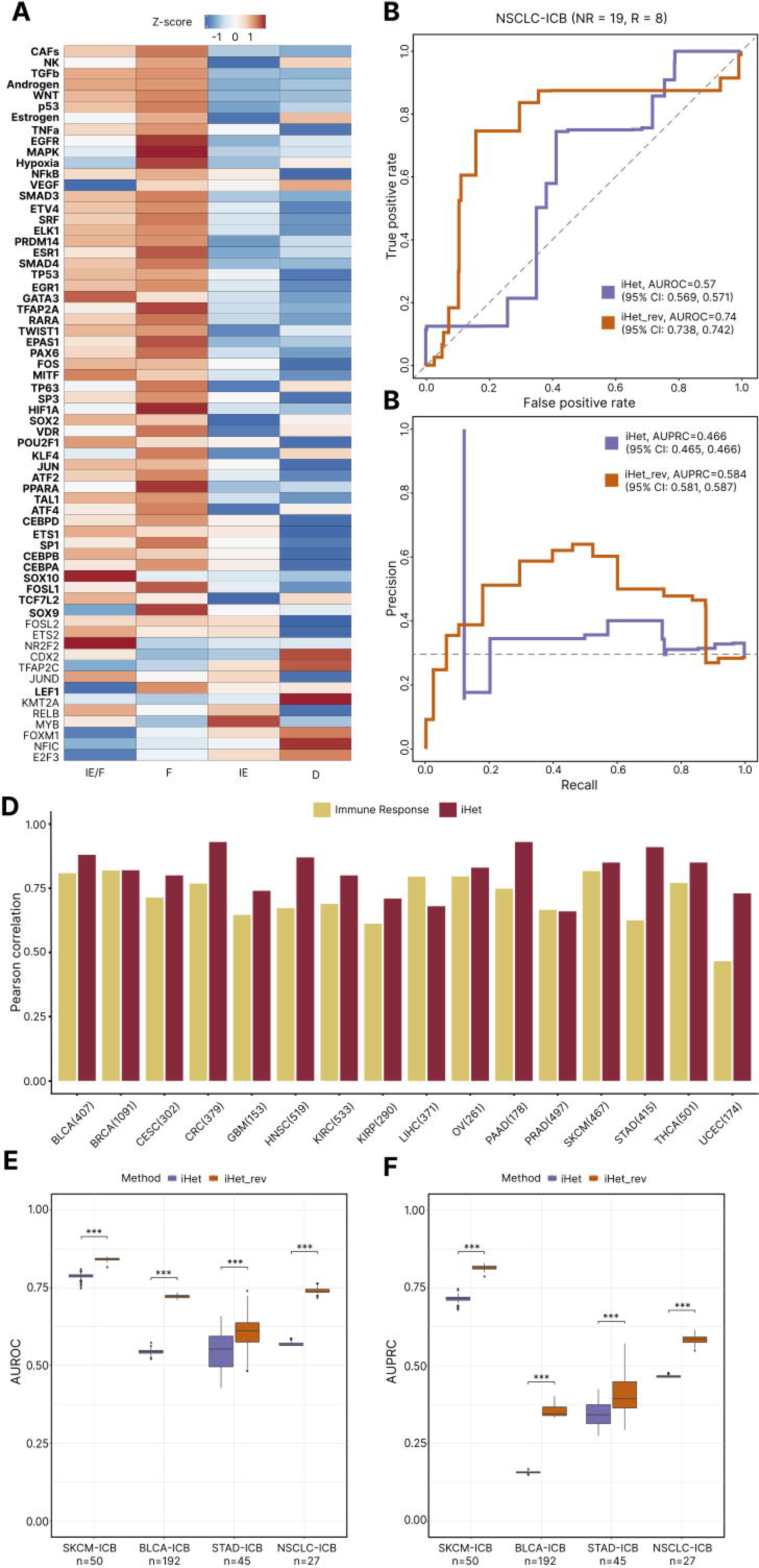
iHet corrected for immune exclusion predicts ICB response in different solid cancers. **(A)** Heatmap showing TCGA-NSCLC patients’ median z-score value of cell fraction, pathway and transcription factor (TF) features (rows) that showed positive correlation (r_s_ > 0, FDR ≤ 0.05) with digital-pathology information about fibroblast density^56^ across different cancer-immune phenotypes (columns). In bold, features that are significantly highly represented (Wilcoxon rank-sum test, FDR ≤ 0.05) in either immune-enriched, fibrotic (IE/F) or fibrotic (F) compared to immune-enriched (IE). Rows (features) were first sorted by modality and then by their effect size across cancer-immune phenotypes. **(B)** Receiving operating characteristic (ROC) curve of ICB response predictions for the NSCLC-ICB cohort (Jung) based on ‘iHet’ and ‘iHet_rev’ scores. ROC curves were computed as the average of the ROC curves obtained for each bootstrap model. **(C)** Precision and recall (PR) curves of ICB response predictions for the NSCLC-ICB cohort (Jung) based on ‘iHet’ and ‘iHet_rev’ scores. PR curves were computed as the average of the PR curves obtained for each bootstrap model. **(D)** Barplots showing the Pearson correlation between the iHet signature of 16 cancer types and both the original iHet signature (TCGA-NSCLC; in gold) and the ensemble immune response score (in garnet). **(E)** Boxplots showing area under the ROC curve (AUROC) values based on ‘iHet’ and ‘iHet_rev’ scores in multiple ICB-cohorts (SKCM: Gide and Auslander, BLCA: Mariathasan, STAD: Kim and NSCLC: Jung). The significance level (∗p < 0.05, ∗∗p < 0.01, ∗∗∗p < 0.001) indicates whether iHet_rev outperforms iHet in terms of ICB response predictions. **(F)** Boxplots showing area under the precision and recall curve (AUPRC) values, based on ‘iHet’ and ‘iHet_rev’ scores in multiple ICB-cohorts (SKCM: Gide and Auslander, BLCA: Mariathasan, STAD: Kim and NSCLC: Jung). The significance level (∗p < 0.05, ∗∗p < 0.01, ∗∗∗p < 0.001) indicates whether iHet_rev outperforms iHet in terms of ICB response predictions. NSCLC represents non-small cell lung cancer, R responders, NR non-responders, AUROC area under the ROC curve, AUPRC area under the precision and recall curve, CI confidence interval, BLCA bladder urothelial carcinoma, BRCA breast invasive carcinoma, CESC cervical and endocervical carcinoma, CRC colorectal adenocarcinoma, GBM glioblastoma multiforme, HNSC head and neck squamous cell carcinoma, KIRC kidney renal clear cell carcinoma, KIRP kidney papillary clear cell carcinoma, LIHC liver hepatocellular, OV ovarian serous cystadenocarcinoma, PAAD pancreatic adenocarcinoma, PRAD prostate adenocarcinoma, SKCM skin cutaneous melanoma, STAD stomach adenocarcinoma, THCA thyroid carcinoma, UCEC uterine corpus endometrial carcinoma.

Given these evidences, we reasoned that disentangling immune exclusion from immune activity can aid the identification of patients which are most likely to respond to immunotherapy with ICB. On this basis, we defined an alternative scoring scheme, called *iHet_rev*, by reversing the signs of the weights of the 64 features associated with immune exclusion. We then calculated both iHet and iHet_rev scores using pre-therapy RNA-seq data generated from a cohort of 27 NSCLC patients treated with anti-PD-1/PD-L1^59^. Compared to the original iHet score, the iHet_rev scoring scheme achieved superior predictive performance in terms of area under the receiver operating characteristic curve (AUROC) (**Figure 3B**, iHet = 0.57 vs. iHet_rev = 0.74). The performance gain was also associated with an increased ability of iHet_rev to distinguish responders from non-responders compared to iHet (**Figure 3C**, area under the precision-recall curve (AUPRC) iHet = 0.47 vs. iHet_rev = 0.58).

To test iHet and iHet_rev more extensively, we considered different cohorts of patients treated with ICB for which both baseline RNA-seq and response data were available, and which have been previously used to test state-of-the-art predictors: patients with skin cutaneous melanoma (SKCM) treated with anti-PD-1 (nivolumab or pembrolizumab)^60,61^, with bladder carcinoma (BLCA) treated with anti-PD-L1 (atezolizumab)^62^, or with and stomach adenocarcinoma (STAD) treated with anti-PD-1 (pembrolizumab)^63^. As these patient cohorts encompass different cancer types, we first assessed whether the iHet signature could be identified in other solid cancers besides NSCLC by running unsupervised MOFA analysis on RNA-seq data from 16 additional TCGA cancer types (n = 6538). This analysis showed that the latent factors F1 identified independently for each cancer type was positively correlated with the TCGA-NSCLC F1 weights (**Figure 3D**, median Pearson correlation between F1 weights r = 0.83), confirming the pan-cancer conservation of the iHet signature. Moreover, the F1 weights across all cancer types were also positively correlated with the predicted immune response score (**Figure 3D and Supplementary Figure 11**, median correlation r = 0.73). This conservation appeared to stem from factors which are essential for the success of immunotherapy, such as the abundance of key antitumor immune cells (e.g. CD8^+^ T cells, M1 macrophages)^64^, intra-cellular signaling pathways and TFs involved in the regulation of PD-L1 and HLA gene expression (NF-κB, TNFα, JAK-STAT, EGFR and TGFβ pathways; NF-κB, RELA, and STAT2 TF)^38,65–67^, and apoptosis (Trail pathway)^68^ (**Supplementary Figure 12**). Differences in feature weights were more evident in cancer types with less marked iHet conservation, like VEGF pathway as well as SPI1 and FOXP1 TFs in prostate adenocarcinoma (PRAD), VEGF pathway in liver hepatocellular carcinoma (LIHC), and CD8^+^ T and T_REG_ cell abundances in glioblastoma (GBM). Given that the iHet signature was conserved in SKCM, BLCA, and STAD, we computed both iHet and iHet_rev scores from the RNA-seq data of all the four ICB cohorts selected above. We compared the performance in terms of both AUROC (**Figure 3E**) and AUPRC (**Figure 3F**) of the iHet and the iHet_rev scores in classifying responders and non-responders, demonstrating that correcting for immune exclusion improves the prediction of ICB response in all four cancer types (p ≤ 0.001). The striking improvement in the BLCA cohort, from 0.55 to 0.70 median AUROC and from 0.16 to 0.34 median AUPRC (baseline AUPRC = 0.15), is in line with the major findings described in the original study, where the lack of response was associated with TGFβ signaling in fibroblasts and exclusion of CD8^+^ T cells from the tumor core^62^.

To better comprehend how the performance of iHet and iHet_rev (**Figure 3E and 3F**) relates to immune infiltration and exclusion, we focused on two cohorts for which a classification of the patients’ cancer-immune phenotypes^8^ was available: the combined Gide and Auslander SKCM cohort^60,61^, with patients classified as “D”, “F”, “IE”, “IE/F”^58^, and the Mariathasan BLCA cohort, with patients classified as immune “desert”, “excluded”, “inflamed”^62^. We divided responding and non-responding patients in tertiles according to iHet or iHet_rev scores, with higher scores being associated with a higher likelihood or response (“High resp”), and vice versa (**Supplementary Figure 13A-B**). For non-responders in both cohorts, iHet_rev resulted in a higher or comparable number of patients assigned to the group with low or medium likelihood of response (“Low resp” and “Mid resp”) and in a lower number of patients assigned to the “High resp” group. As expected, these changes were primarily due to reclassification of fibrotic/excluded samples, with a marked reduction in the BCLA cohort of excluded and desert phenotypes in the “High resp” group of non-responders (**Supplementary Figure 13A)**. Among responders, iHet_rev resulted in an increased number of patients in the “High resp” group, especially in the BLCA cohort. In iHet_rev compared to iHet, the fraction of responders increased from 11/20 to 12/20 and from 8/23 to 14/23 in the SKCM and BLCA cohort, respectively. Compared to iHet, iHet_rev showed improved predictive performance of ICB response, suggesting a better discrimination of immune activity from fibroblast-mediated immune exclusion.

These results show that the iHet score computed from pre-therapy tumor RNA-seq data predicts patients’ response to ICB in different solid cancers. Most importantly, by integrating orthogonal information from digital pathology data, we could disentangle concomitant but opposing mechanisms underlying immune activity versus immune exclusion, further augmenting the predictive value of iHet for ICB immunotherapy.

### Integration of tumor mutational burden improves iHet predictive value for ICB immunotherapy

By disentangling the features associated with immune exclusion, we could improve the predictive value of iHet for ICB immunotherapy. However, as its predictions were still imperfect, we considered orthogonal drivers of therapeutic response that might be not captured by the cellular and molecular representation of anticancer immune responses provided by iHet. Apart from the composition and crosstalk in the TME, a complementary hallmark of response to ICB is the tumor “foreignness”, namely the potential of tumor cells to present neoantigens that are recognized as “non self” by the immune system^54,69^. We quantified tumor foreignness using tumor mutational burden (TMB) for all the cohorts for which this information could be accessed or computed (**STAR Methods**), and tested its association with the iHet scores (**Supplementary Figure 14A**). iHet scores showed negative or no correlation with TMB in all datasets except TCGA breast cancer (BRCA), colorectal cancer (CRC), and kidney renal clear cell carcinoma (KIRC), with a particularly strong, negative correlation in the multi-biopsy data (r = -0.46 and -0.60 for Jia and Sharma, respectively). By deriving iHet scores from TRACERx data using the two alternative approaches described previously (i.e., unsupervised MOFA analysis and matrix multiplication based on the TCGA-NSCLC weights), we could confirm no association of iHet with TMB also in this validation dataset (**Supplementary** Figure 14B).

Given the orthogonal information provided by TMB, as suggested by these results, we decided to consider it for all patient cohorts for which this information was available, and assess whether we could use it to improve iHet predictive performance. Specifically, we quantified and integrated three scores representing major determinants of patients’ response to ICB: 1) anticancer immune response, quantified via iHet; 2) immune exclusion, quantified using all the features associated with immune exclusion (iHet_excl) (**Supplementary Table 2**), but considered with a negative weight due to their negative impact on response (**STAR Methods**); and 3) TMB as a proxy for tumor foreignness. We observed that a weighted combination of the three scores can improve the predictive performance with respect to each individual score, although the role played by each individual component was dependent on the specific cohort (**Figure 4**). Importantly, combining all the three components with equal weights provided better predictions than each individual score in BLCA and NSCLC cohorts, confirming the importance of integrating different hallmarks of response to ICB therapy.

**Figure 4.**
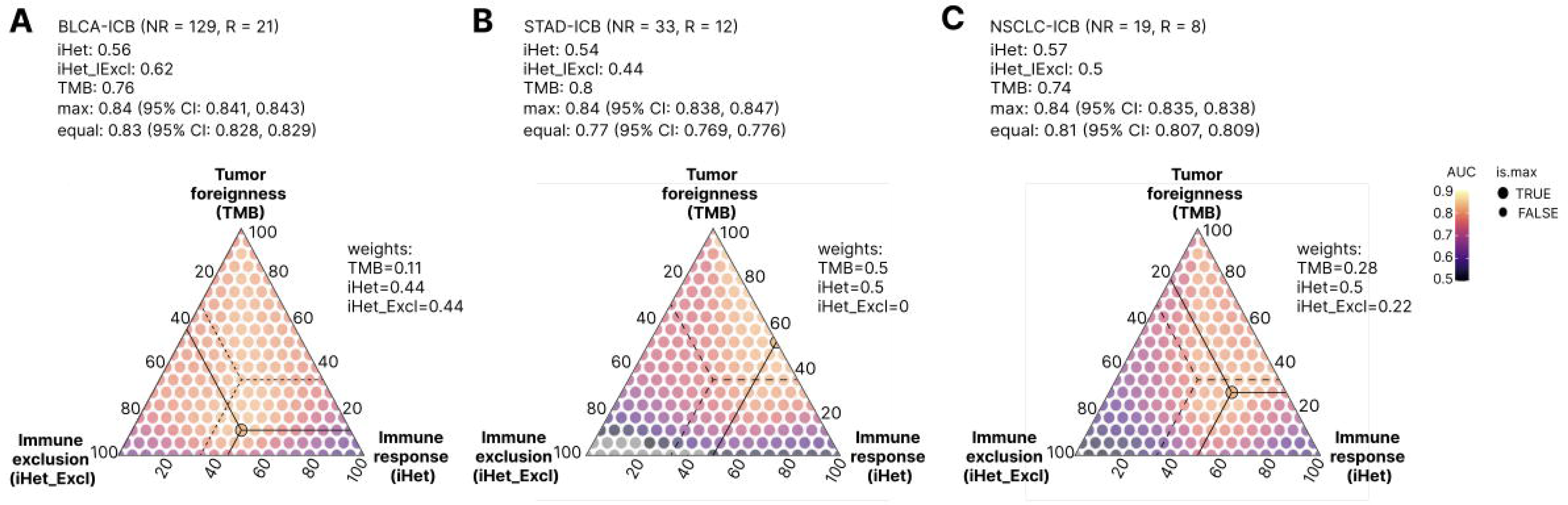
Integration of different hallmarks of anticancer immunity improves the prediction of ICB response. Predictive performance for three ICB-cohorts: **(A)** bladder urothelial carcinoma (BLCA, Mariathasan cohort), **(B)** stomach adenocarcinoma (STAD, Kim cohort), and **(C)** non-small cell lung cancer (NSCLC, Jung cohort). The triangle plots show the area under the ROC curve (AUC) obtained using an ensemble predictor computed as a weighted average of scores of: 1. immune response (‘iHet’), 2. immune exclusion (‘iHet_excl’) and 3. tumor foreignness (TMB). Best performances (max) are highlighted with a circle and continuous lines are projections to the corresponding weights showing the percentage contribution of the different scores. The vertices correspond to individual contributions of each score. The center of the triangle corresponds to the performances (equal) achieved with equal contributions of the three scores (projections to the weights are shown using dotted lines). CI: confidence interval.

Taken together, these results show that the integration of multimodal data capturing different facets of the TME shed light on the mechanisms that contribute to determining the success or failure of anticancer immune immunity, and can be used to predict patients’ response to ICB.

## Discussion

ICB immunotherapy has revolutionized oncology and our understanding of anticancer immunity, further unveiling a crucial role of the TME for determining disease progression and response to therapy. The successes and failures that have punctuated the immuno-oncology field in the past decade has shown the path to redesign the original “precision oncology” paradigm: enlarging the center of attention from the sole genetic makeup of malignant cells to a much wider and intricate crosstalk taking place among all cells of the TME^70^. RNA-seq technologies can generate rich datasets capturing the molecular complexity of the TME and characterize the components of inter- and intra-tumor phenotypic diversity that ultimately impact patients’ responses to (immuno)therapy.

To date, most of the attempts to anticipate patients’ responses to ICB immunotherapy from the analysis of pre-therapy, tumor RNA-seq data have remained entangled to a gene-centric implementation^14–16,30–33^, which risks to derive partially redundant gene signatures with limited potential for discovering the mechanistic basis of anticancer immune responses. In this study, we took a *systems* approach, using RNA-seq data from more than thousand patients with NSCLC to extract a holistic view onto the cell types, pathways, and transcription factors underlying intra- and inter-tumor heterogeneity. Through unsupervised analysis of this multimodal data, we pinpointed a latent factor capturing most of the data heterogeneity. This factor was identified in different NSCLC cohorts but not in healthy-lung RNA-seq data, suggesting its relevance in a disease context. Through the independent analysis of additional RNA-seq data from more than 6000 TCGA patients we demonstrated that this latent factor is conserved in different solid cancers. Of note, unlike our previous effort focused on the supervised modeling of anticancer immune response^29^, this factor was derived in a completely *unsupervised* manner. Nevertheless, we could demonstrate its strong association with the predicted immune response in all the analyzed cancer types. This result underscores once more the central role of the non-malignant component of the TME in determining the fate of anticancer immune responses. Given its association with TME heterogeneity and anticancer immunity, we termed this latent factor *immune heterogeneity (iHet)* signature.

A major strength of our systems approach lies in its direct interpretability. For instance, the top features positively associated with iHet reflect the infiltration of T lymphocytes (CD8^+^ T cells, CD4^+^ T cells, and T_REG_ cells), and the response to IFN-γ signaling (JAK-STAT pathway, STAT1 and STAT2 transcription factors), which ratifies the positive association of the iHet signature with immune response. CD8^+^ T-cell abundances were not always consistent across biopsies collected from the same patient, suggesting a spatial, intra-tumoral variation of T-cell infiltration in some NSCLC tumors. One limitation of the present work is that we did not consider DC in our bulk analysis. We disregarded this cell type for the difficulty to quantify them via deconvolution from bulk RNA-seq data^71^ due to their low abundance and broad transcriptional diversity. Nevertheless, DC are critical players in the orchestration of anticancer immune responses, acting not only in draining lymph nodes but also in situ in the TME^11^.

Thanks to the curation of a single-cell atlas of the NSCLC TME and its characterization in terms of “systems” features rather than gene expression, we could trace back the major molecular programs identified in bulk RNA-seq data to their cellular origin. Our results from the bulk and single-cell data suggest a major contribution of myeloid cells to the heterogeneity of NSCLC TME. SP1 (PU.1) transcription factor was the feature with the largest positive weight both in the TCGA and multi-biopsy RNA-seq data; according to single-cell data, its high activity is a prerogative of myeloid cells, in accordance with its role as master regulator of myeloid-cell differentiation^72^. Among the top iHet features associated with immune response are other myeloid cell-enriched transcription factors and pathways. Some of these features presented preferential activity in specific subsets, including MYCN (TAM and alveolar macrophages) and RFX5 (cDC1, cDC2, and “langerhans-like” cD1A^+^ cDC2) transcription factors. NF-κB and TNFα pathways showed activity across most of myeloid cells, but were particularly enriched in classical monocytes. TNFα and NF-κB were also the features showing the largest intra-tumor variability, suggesting the spatial segregation of myeloid cells in the TME.

Among the top features negatively associated with iHet were the E2F4 and TFDP1 cell-cycle regulators. Our analysis of single-cell data showed that these transcription factors had a higher activity not only in small subsets of cycling cells across different lineages, as expected, but also in a large proportion of tumor cells. This result aligns with the conserved programmes underlying the phenotypic heterogeneity and plasticity of tumor cells identified in several pan-cancer single-cell studies of tumor samples and cancer cell lines^73–75^.

Interestingly, some of the top-features positively associated with iHet underlie mechanisms of immune exclusion rather than anticancer immunity, like the presence of T_REG_ cells and CAFs, and the activity of the TGFβ pathway. These immuno-suppressive features may result associated with immune infiltration due to their co-occurrence in bulk-tumor RNA-seq data with other immune response mechanisms. A prototypical example of such “spurious” correlations detected in bulk tumors are T_REG_ cells, whose presence has been associated with a good clinical outcome in some cancer patients likely due to their positive association with overall T-cell infiltration^36,76^. Here, we focused on the immuno-suppressive role of CAFs, a type of fibroblasts that are able to form collagen-rich fibrotic structures that restrict T-cell infiltration into the core of the tumor^62,77^. Through regulation of TGFβ signaling, CAFs can also promote T_REG_ cell differentiation^78^ and hamper the expansion of T stem-like cells^79,80^. By integrating orthogonal information from digital pathology data, we could identify the iHet features associated with the density of fibroblasts in tumors, thereby disentangling immune activity from immune exclusion mechanisms. The top associated features included not only CAF abundances derived via deconvolution, but also the activity of TGFβ pathway and SMAD3/SMAD4 transcription factors, in agreement with their role as master regulators of tissue fibrosis^57^. Of note, this approach also identified some features that might not be direct drivers or mediators of immune exclusion, but rather co-occurring mechanisms that cannot be disentangled from fibrosis using bulk RNA-seq data.

Intrigued by the strong signal associated with myeloid cells and fibroblasts in our bulk and single-cell analyses, we further characterized the spatial organization of these two cell lineages using lung cancer spatial transcriptomics data. Myeloid cells and fibroblasts are two major components of the TME that interact with each other and influence tumor progression and response to therapy. Characterized by an extreme plasticity, they can play both pro- and anti-tumorigenic roles and are the object of intense research to identify therapeutic targets for their modulation or reprogramming^81–83^. Our analysis of NSCLC spatial transcriptomics, despite limited by the current data availability, confirmed the high abundance and geospatial heterogeneity of fibroblasts and myeloid cells, and confirmed peribronchial fibroblasts and classical monocytes as the major sources of TGFβ and TNFα/NF-κB pathway activity, respectively.

Through the analysis of pre-therapy RNA-seq data from 272 patients treated with ICB we demonstrated that iHet, despite being derived with an unsupervised approach, predicts patients’ response to ICB immunotherapy. Most notably, by disentangling the immune exclusion component, we could derive a scoring scheme (iHet_rev) with improved prediction performance. This is in line with preclinical studies suggesting that resistance to ICB is partly determined by TGFβ signaling^77,80,84^. Incorporating TMB information further increased the prediction performance confirming that it constitutes a complementary hallmark of anticancer immunity, not necessarily correlated with immune infiltration and activity in bulk tumors^85,86^. The negative association between iHet and TMB, seen only in the multi-biopsy data, may reflect tumor immunoediting^87^ but could also result from the difficulty to reliably detect somatic mutations in tumors with higher immune infiltration –and lower tumor purity– which could be exacerbated in small biopsy samples.

The value of the iHet lies in its capability to provide an holistic, quantitative portrait of TME heterogeneity which is easily interpretable. Although deriving a predictive score was not the overarching aim of this work, the high predictive value of this approach underscores its value for cancer immunology and immunotherapy, and the importance of modeling the different facets of tumor heterogeneity. It is likely that the prediction of patients’ response to ICB can be further improved by incorporating additional drivers of anticancer immune responses and mechanisms of immune evasion, like the presence of tertiary lymphoid structures^15,88^, antigen presentation defects^89,90^, neoantigen burden and “quality”^91–93^, as well as cancer and germline genetics^94^.

Our analysis of NSCLC data showed that inter-patient heterogeneity is more prominent than intra-patient one. Nevertheless, the analysis of multi-biopsy and spatial transcriptomics data revealed regional variability of immune and stromal cells, together with their molecular underpinnings, which were more marked in some tumors. Notably, intra-sample phenotypic heterogeneity also had an impact on the iHet scores computed from each biopsy, which was extensive in some tumors and more limited in others. Transcriptomic intra-tumor heterogeneity is a major confounder of expression-based biomarkers^17^ and can represent a major hurdle for clinical decision making. In the context of therapy response prediction, this spatial variability strongly limits our capability to anticipate patients’ trajectories from small biopsies. Similarly, the coexistence of different cancer-immune phenotypes in the same tumor complicates the prediction of how the TME will evolve and which force will prevail, ultimately determining the clinical outcome. In this light, the availability of new technologies for the generation of spatially-resolved omics data^95–97^ can help us shed light on local cell-cell interaction mechanisms that shape the evolution of the TME in space and time.

## Acknowledgements

FF was supported by the Austrian Science Fund (FWF) (no. T 974-B30 and FG 2500-B) and by the Oesterreichische Nationalbank (OeNB) (no. 18496). GS was supported by a DOC fellowship of the Austrian Academy of Sciences. ZT was supported by the European Research Council (ERC) under the European Union’s Horizon 2020 Research and Innovation Programme (grant agreement no. 786295). NFCCdM was supported by the European Research Council (ERC) under the European Union’s Horizon 2020 Research and Innovation Programme (grant agreement no. 852832). NM is a Sir Henry Dale Fellow, jointly funded by the Wellcome Trust and the Royal Society (Grant Number 211179/Z/18/Z), and also receives funding from Cancer Research UK Lung Cancer Centre of Excellence, Rosetrees, and the NIHR BRC at University College London Hospitals. The computational results presented here have been achieved in part using the LEO HPC infrastructure of the University of Innsbruck. The results shown here are in part based upon data generated by TCGA Research Network (https://cancergenome.nih.gov) and by the TRACERx study (http://tracerx.co.uk/). The authors would like to thank Dr. Ricard Argelaguet for providing useful suggestions for setting up the multimodal data analysis.

## Author contributions

Conceptualization: FF, FE, OLS; Methodology: FF, FE, OLS, GS; Software: GS, OLS, CZ, MZ, DR; Formal analysis: GS, OLS, JK, CZ, MZ, MA, NM, DR; Investigation: FF, FE, JK, GS, OLS, NFdCCdM; Resources: FF, DR, ZT; Data Curation: GS, OLS, CZ; Writing - Original draft: FF, FE, OLS; Writing - Review & Editing: all; Visualization: GS, OLS, CZ; Supervision: FF, FE; Project Administration: FF.

## Declaration of interests

FF consults for iOnctura. FE is a freelancer for TheraMe! AG. GS is an employee of Boehringer Ingelheim International Pharma GmbH & Co KG, Biberach, Germany. NM has received consultancy fees and has stock options in Achilles Therapeutics. NM holds European patents relating to targeting neoantigens (PCT/EP2016/ 059401), identifying patient response to immune checkpoint blockade (PCT/ EP2016/071471), determining HLA LOH (PCT/GB2018/052004), predicting survival rates of patients with cancer (PCT/GB2020/050221).

## Data and code availability

- No new data was generated for this study. An overview of the bulk sequencing datasets used (including accession numbers) is reported in **Supplementary Table 1**.
- The spatial transcriptomics dataset was obtained from the 10x Genomics website: https://www.10xgenomics.com/resources/datasets/human-lung-cancer-ffpe-2-standard.
- The single-cell lung cancer atlas, version 2022.05.10, was obtained from Zenodo: https://doi.org/10.5281/zenodo.7104046
- Processed input data, as well as intermediate and final results of the computational analysis are available from FigShare: https://doi.org/10.6084/m9.figshare.24855882
- The source code required to reproduce the Analyses is available from GitHub: https://github.com/ComputationalBiomedicineGroup/iHet.

## STAR Methods

### Bulk RNA and whole-exome sequencing data access and preprocessing

Preprocessed RNA-seq data for 18 TCGA solid tumors, including LUAD and LUSC, were downloaded via the Broad GDAC Firehose (https://gdac.broadinstitute.org), released January 28, 2016. We selected only samples from primary or, for melanoma, metastatic tumors from 7,750 patients. Gene expression data were extracted from the “illuminahiseq_rnaseqv2-RSEM_genes” files as raw counts, corresponding to the “raw_count” values, and as TPM, computed by multiplying the “scaled_estimate” values by 1e6. Genes with non-valid HGNC symbols were removed, and gene expression levels across identical HGNC symbols were averaged.

Preprocessed RNA-seq data from the GTEx project (v8 release)^98^ in the form of gene counts and TPM were obtained from the GTEx portal (https://www.gtexportal.org/home/). We selected only lung samples and averaged expression levels across genes with identical HGNC symbols. Raw RNA-seq and WES data (FASTQ files) from the Jia et al.^18^ and Sharma et al.^99^ studies, as well as for four cohorts of cancer patients treated with ICBs (Gide, Auslander, Kim, and Jung) were downloaded from the Sequence Read Archive (SRA, https://www.ncbi.nlm.nih.gov/sra). Raw RNA-seq and WES data from the TRACERx study^17^ and the Mariathasan cohort^62^ were available on controlled access through the European Genome-Phenome Archive (EGA, https://ega-archive.org/).

An overview of the analyzed bulk sequencing datasets, together with their accession numbers, is provided in **Supplementary Table 1.**

quanTIseq was used to process the RNA-seq FASTQ files and obtain gene counts and TPM as done previously^21^. Briefly, raw RNA-seq reads were pre-processed with Trimmomatic^100^ to remove adapter sequences and read ends with Phred quality scores lower than 20, discard reads shorter than 36 bp, and trim long reads to a maximum length of 50 bp. The pre-processed RNA-seq reads were analyzed with Kallisto^101^ to generate gene counts and transcripts per millions (TPM) using the “hg19_M_rCRS” human reference.

### Computation of tumor mutational burden

Tumor mutational burden (TMB) estimates for TCGA patients were available from “The Immune Landscape of Cancer”^102^ data repository on the Genomics Data Commons (GDC) portal (“mutation-load_updated.txt” file). TMB was calculated as the sum of silent and non-silent mutations per mega-base (Mb).

Jia and Sharma raw sequencing data were analyzed with NextNEOpi^92^ to derive TMB estimates as the number of somatic mutations per megabase. Briefly, tumor and normal paired-end FASTQ files (from RNA and DNA sequencing) were analyzed with NextNEOpi (version 1.2, default parameter settings, Nextflow^103^ version 21.10.0) providing sex information for each individual, wherever available (i.e. “XX” or “XY”).

The TMB per Mb for each tumor region within the TRACERx cohort was calculated following the guidelines outlined in the Friends of Cancer Research TMB Harmonization Project^104^. In brief, based on a bed file described in the associated publication, we considered any non-silent coding somatic mutations within the 32.102 Mb of the genome considered exonic. TMB/Mb was then calculated for each tumor region by simply calculating the number of mutations divided by 32.102, to yield an estimate of mutations per Mb.

For Jung, Kim, and Mariathasan ICB cohorts, we considered the available measurements of TMB. In the Jung and Kim cohorts, the TMB was quantified as the total number of non-synonymous mutations derived from WES data. In the Mariathasan cohort, this was defined as the total number of both synonymous and non-synonymous mutations derived from panel sequencing. The TMB was provided as mutations per megabase, except for the Kim cohort, for which the TMB was available as a three-level categorical variable: low (< 100), moderate (100 – 400), and high (> 400).

### Quantification of immune-cell fractions, pathway, and transcription-factor activities from bulk RNA sequencing data

We used quanTIseq deconvolution method^21^ to quantify from RNA-seq TPM data cell fractions for ten different immune cell types: B cells, classically (M1) and alternatively (M2) activated macrophages, monocytes, neutrophils, natural killer cells, non-regulatory CD4^+^ T cells, CD8^+^ T cells, regulatory T (T_reg_) cells, and myeloid dendritic cells (DC). quanTIseq was run with default parameter settings, except “tumor = TRUE” and “rawcounts = TRUE”. As non-regulatory CD4^+^ T cells are difficult to estimate via deconvolution, we considered instead total CD4^+^ T cells computed as the sum of non-regulatory and regulatory (T_reg_) cells. Due to the difficulty of estimating DC from bulk RNA-seq data^71^, which results in sparse and noisy cell fractions, we discarded this cell type from downstream analyses. In addition to immune cells, we quantified the fractions of two additional cell types playing a major role in the TME, cancer associated fibroblasts (CAFs) and endothelial cells, estimated with EPIC deconvolution method^22^ accessed through the immunedeconv R package^71^ (with “tumor = TRUE” and “scale_mrna = TRUE” parameter settings).

RNA-seq gene counts normalized with variance stabilizing transformation (VST) implemented in DESeq2^105^ were analyzed with PROGENy to quantify the activity of 14 pathways: Androgen, EGFR, Estrogen, Hypoxia, JAK-STAT, MAPK, NF-κB, p53, PI3K, TGFβ, TNFα, Trail, VEGF and WNT^23,106^. The top 100 target genes of the progeny model were used. TPM gene counts were analyzed with DoRothEA^24^ to compute TF activity scores, considering only high-confidence regulons (class “A” or “B”). PROGENy and DoRothEA analyses were performed using the EaSIeR R package^29^ (with “remove.genes.ICB_proxies = FALSE” settings).

### Multi-Omics Factor Analysis (MOFA)

Cell type fractions were first converted to percentages and then log10-transformed (pseudo-count = 0.001), whereas pathway activities were scaled to have unit variance. The range of transcription factor activities was already comparable (these were inferred using Viper^107^ which returns a normalized enrichment score). All data modalities had a gaussian-like distribution, consistent with MOFA’s modeling assumption.

We exploited MOFA’s multi-group functionality to fit models for TCGA-NSCLC and Jia-Sharma using LUAD and LUSC and Jia and Sharma as separate groups, respectively. This allowed us to identify sources of variability shared between groups but also exclusive for each group (i.e. shared factor weights across groups). In this case, cell type, pathway and transcription factor features were centered per group. We trained MOFA models with 7 factors to be optimized, “medium” convergence mode, and max. 7000 iterations and otherwise default parameters. MOFA models were learned separately for each dataset. Model training was repeated for 100 randomly generated bootstrap samples.

### Computation of the immune response score from bulk RNA sequencing data

Starting from gene TPM, we computed an ensemble immune response score by integrating nine state-of-the-art predictors of anticancer responses^14–16,30–33^ that were previously shown to have high agreement across different cancer types^29^. These immune response scores where computed via the ‘compute_gold_standards’ function of the EaSIeR R package^29^, specifying the following ‘list_gold_standards’: “CYT”, “Roh_IS”, “chemokines”, “Davoli_IS”, “IFNy”, “Ayers_expIS”, “Tcell_inflamed”, “RIR”, “TLS”. The scores were z-scored considering their mean and standard deviation across the entire TCGA pan-cancer cohort, and mediated to obtain a single immune response score per sample.

### Assessing inter vs. intra patient heterogeneity

We employed two metrics for assessing the inter- and intra-patient heterogeneity of each feature in datasets with multiple biopsies from the same patient (Jia, Sharma, and TRACERx): (1) a “feature stability-score” based on kendall-rank correlation and (2) the ratio of median absolute deviation (MAD) of the patient-medians vs. the median of the MADs within each patient. The feature stability score was computed as follows: Over 100 iterations, for each patient, a random sample was chosen with replacement. Then, for each feature, a correlation matrix was calculated between the 100 random samples using the Kendall rank coefficient. Finally, the stability score was obtained as the median of the upper triangle of the correlation matrix.

### Building a single-cell atlas for studying the TME

We previously published the single-cell lung cancer atlas (LuCA)^39^ comprising >1.2M cells from 318 patients and 29 datasets from both healthy lung donors, COPD and cancer patients measured on 6 different sequencing platforms. In summary, in that study, the individual scRNA-seq datasets were quality controlled and low-quality cells were filtered based on dataset-specific thresholds for the number of detected genes, UMI counts, and the fraction of mitochondrial reads. Then, datasets were merged, gene identifiers harmonized and a batch effect-corrected latent embedding computed using scANVI^50^. Doublets were removed using the SOLO algorithm^108^ and cell types annotated based on unsupervised leiden clustering^109^ and a curated list of marker genes. Finally, two additional datasets were added to the LuCA “core” atlas by projecting them on the annotated data using scArches^110^ and their cell-type labels inferred automatically based on a nearest neighbor graph majority voting.

Since the focus of this study is on cancer patients, we created a subset of LuCA including only primary tumor and normal adjacent tissues of cancer patients of LUAD and LUSC histology, containing 801,488 cells from 208 patients. Neighborhood graph and UMAP embedding^111^ were recomputed on the subset based on the original scANVI latent representation. The subset used in this study is based on the 2022.05.10 version^112^ of LuCA.

### Cell-type annotation

While LuCA provides annotations for 44 cell types, the annotation of the myeloid cluster is relatively coarse-grained. We, therefore, refined the annotation of myeloid subsets based on unsupervised clustering and known marker genes. Clusters were computed using the Leiden algorithm^109^ as implemented in scanpy^113^ based on a neighborhood graph obtained from the scANVI embeddings. We assigned two Macrophage subsets, three DC subsets and two monocyte subsets as well as clusters of cycling Macrophages, Monocytes and DCs, respectively (see **Supplementary Figures 8a-b**). All clusters were backed by multiple patients and datasets (**Supplementary Figure 8c**).

### Functional analyses of scRNA-seq data

We performed pathway and transcription factor (TF) signaling analysis on primary tumor samples with PROGENy^23,51^ and DoROthEA^24,51^, respectively. Scores were computed using the *dorothea-py* and *progeny-py* packages. The top 1,000 target genes of the progeny model were used, as recommended for single-cell data. For dorothea, only regulons of the highest confidence levels “A” and “B” were used. The methods were run with the options *num_perm=0*, *center=True*, *norm=True, scale=True*, and *min_size=5*.

### Analysis of spatial transcriptomics data

Preprocessed counts generated with the 10x Genomics (www.10xgenomics.com/) spatial transcriptomics technology from a lung squamous cell carcinoma tissue section were downloaded from the 10x Genomics website (“Human Lung Cancer (FFPE)” dataset). To perform spatial cell-type deconvolution, we used the Lambrechts study^42^ as reference dataset, as contained in our NSCLC single-cell atlas. The Lambrechts subset was chosen for the technology used to generate the data (i.e. 10x) and for the high cell-type coverage. From this reference dataset cycling cell types were removed, resulting in a total of 38,312 cells.

Spatial analysis was performed using the R package spacedeconv (https://github.com/omnideconv/spacedeconv). Both the single-cell reference dataset and the spatial slide were preprocessed with the ‘preprocess’ function of the spacedeconv package with default settings, removing observations (cells or spots) with a unique molecular identifier (UMI) count below the default threshold of 500 and eliminating genes with zero counts across all observations.

We used the spacedeconv package to apply the cell2location^52^ deconvolution method to spatial transcriptomic data to reconstruct the spatial organization of the different cell types composing the TME. Cell2location was run with raw counts for both spatial and single-cell reference data. Cell2location operates in a two-step approach, including scRNA-seq-informed signature building followed by deconvolution of the spatial transcriptomics object. To build the signature we used the ‘build_model’ function of spacedeconv with the following settings: epochs = 250 (number of epochs to train the cell2location model), gpu = TRUE (whether to train on GPU), and assay_sc = “counts” (assay to extract from the single-cell object analyzed). For the second step, we used the ‘deconvolutè function of spacedeconv with the following parameters: epochs = 30000, gpu=TRUE, assay_sp = “counts” (assay to extract from the spatial transcriptomics object analyzed).

For pathway activity analysis, spatial data was normalized as counts per million (CPM) and analyzed with decoupleR^53^ within the spacedeconv framework. We accessed PROGENy^23^ annotations using the ‘get_decoupleR_referencè function with default parameters in spacedeconv. Subsequently, we employed the ‘compute_decoupleR_activities’ function with parameters ref = “wmean” and assay = “cpm”.

The results from the cell-type deconvolution and pathway activity inference were visualized using spacedeconv ‘plot_celltypè function with default settings. To create the “comparative” plots of the monocyte-fibroblast interface, we computed the ratio of fibroblasts vs. monocyte cell fractions, adding a pseudocount of 1 to avoid division by zero. A similar approach was used for pathway scores but, as they are adimensional and, thus, not directly comparable, we rescaled them in the [1,2] range before computing the ratio. The resulting ratios were then log-scaled.

### Definition of cancer-immune phenotypes

We used the four cancer-immune phenotypes defined by Bagaev et al.^58^ from molecular data: immune-enriched fibrotic (IE/F), immune-enriched (IE), fibrotic (F) and immune-depleted (D). These were available for 989 TCGA-NSCLC patients and melanoma patients from Gide and Auslander cohorts. For patients from the Mariathasan BLCA cohort, we used an available classification of three cancer-immune subtypes based on CD8^+^ T-cell immunohistochemistry: immune inflamed, immune excluded and immune desert.

### Association of iHet scores with fibroblast density

We used the feature “DENSITY [FIBROBLAST CELLS] in [TUMOR]_HE” for TCGA-NSCLC patients, as provided by the original work^56^. This was automatically extracted from H&E pathology slides. We first assessed which iHet features were positively correlated (Spearman correlation r > 0, FDR ≤ 0.05) with this information about the spatial organization of fibroblasts (see **Supplementary Table 4**), and then evaluated the probability of iHet features to be greater in fibrotic than in immune-enriched tissues (Wilcoxon rank-sum test, FDR ≤ 0.05).

### Computation of iHet, iHet_rev, and iHet_excl

First, features were computed and normalized using the mean and standard deviation from TCGA MOFA bootstrap models. More specifically, we subtracted the TCGA mean from all descriptors and additionally divided the pathway activity scores by the TCGA standard deviation. For each bootstrap model, the iHet score for each patient was computed by a matrix multiplication between the weights of iHet (from MOFA analysis) and the values of the normalized features.

To calculate the iHet_rev score, we first reversed the sign of the MOFA weights that were positively correlated with the density of fibroblasts in tumor region (r_s_ > 0, FDR ≤ 0.05) and then applied the matrix multiplication between the weights and the normalized features. Similarly, the iHet_excl score was defined by inverting the score resulting from the matrix multiplication of only the subset of the features that showed positive correlation.

### Classification of response to checkpoint blockers inhibitors

We only considered patients treated with anti-PD-L1 or anti-PD-1. For the Jung dataset, we considered responders and non-responders as “patients with durable clinical benefit” and “non-durable clinical benefit”, respectively. For the Gide, Mariathasan and Kim datasets we considered responders as patients with “complete response” or “partial response”, and non-responder as patients with “progressive disease” or “stable disease”. For the Auslander cohort we used the available definition of responders and non-responders, as response evaluation criteria in solid tumors (RECIST) were not provided.

### Weighted combination of iHet, iHet_excl and TMB

For the Kim-ICB cohort we used the classification of low TMB (TMB*_L_*), moderate TMB (TMB*_M_*), and high TMB (TMB*_H_*) as provided in Kim et al.^63^. For Jung and Mariathasan-ICB cohorts, we grouped patients in thirds as described in Carbone et al.^114^: lower (TMB*_L_*), intermediate (TMB*_M_*) and upper (TMB*_H_*) tertile.

The final prediction for each patient *i* was obtained by computing the weighted average between the iHet, the iHet_excl and the TMB scores (MOFA feature weights to compute iHet and iHet_excl scores can be found in **Supplementary Table 2**). Scores were first scaled between 0 and 1 to make them comparable, such that TMB*_L_* = 0, TMB*_M_* = 0.5, and TMB*_H_* = 1. The relative weight is given by the hyperparameter η as described in the following equation:

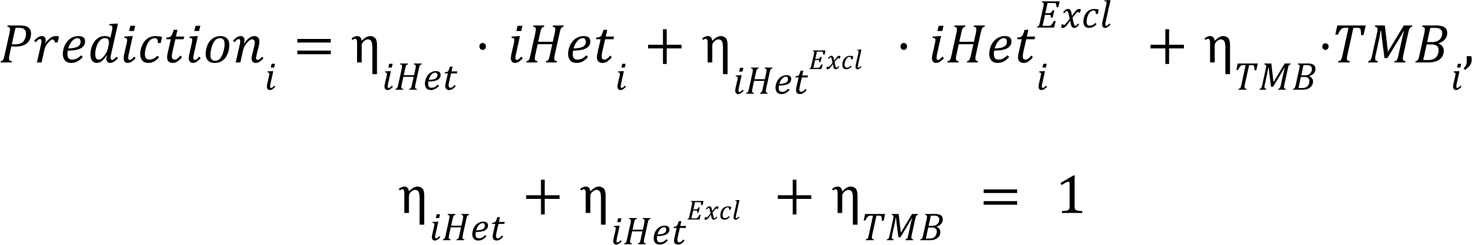

## Supplementary Tables

**Supplementary Table 1.** High-throughput sequencing datasets analyzed in this study.

**Supplementary Table 2.** Median feature weights of iHet, iHet_rev, and iHet_excl.

**Supplementary Table 3.** Average PROGENy pathway scores and Dorothea transcription factor scores per cell type estimated from single-cell NSCLC atlas.

**Supplementary Table 4.** Spearman correlation between iHet features and fibroblast density estimated from digital pathology tumor images.

## Notes

https://github.com/ComputationalBiomedicineGroup/iHet

