## Supplementary Figures 1-14 for "Multimodal analysis unveils tumor microenvironment heterogeneity linked to immune activity and evasion"

### SUPPLEMENTAL FIGURES

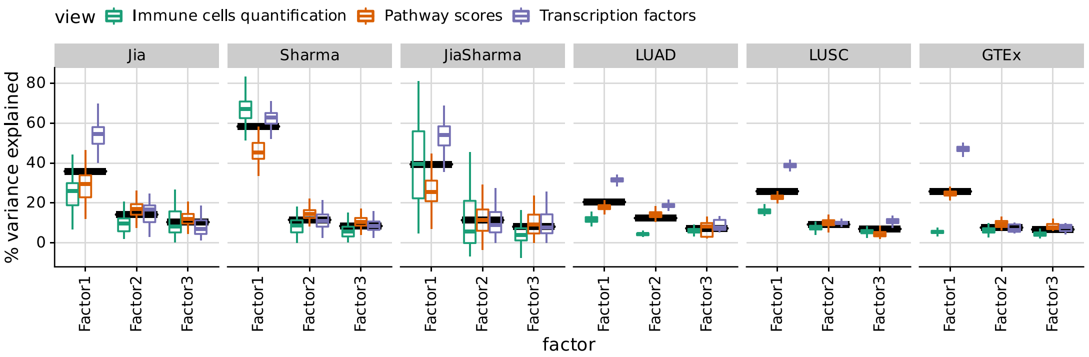


**Figure S1. Explained variance per factor according to MOFA across 100 bootstrap runs.** Of the boxplots, the central line denotes the median. The boxes extend to the 25 and 75 percentiles. The whiskers extend to the most extreme data point within 1.5 of the interquartile range (IQR). The black bar denotes the mean of all values.

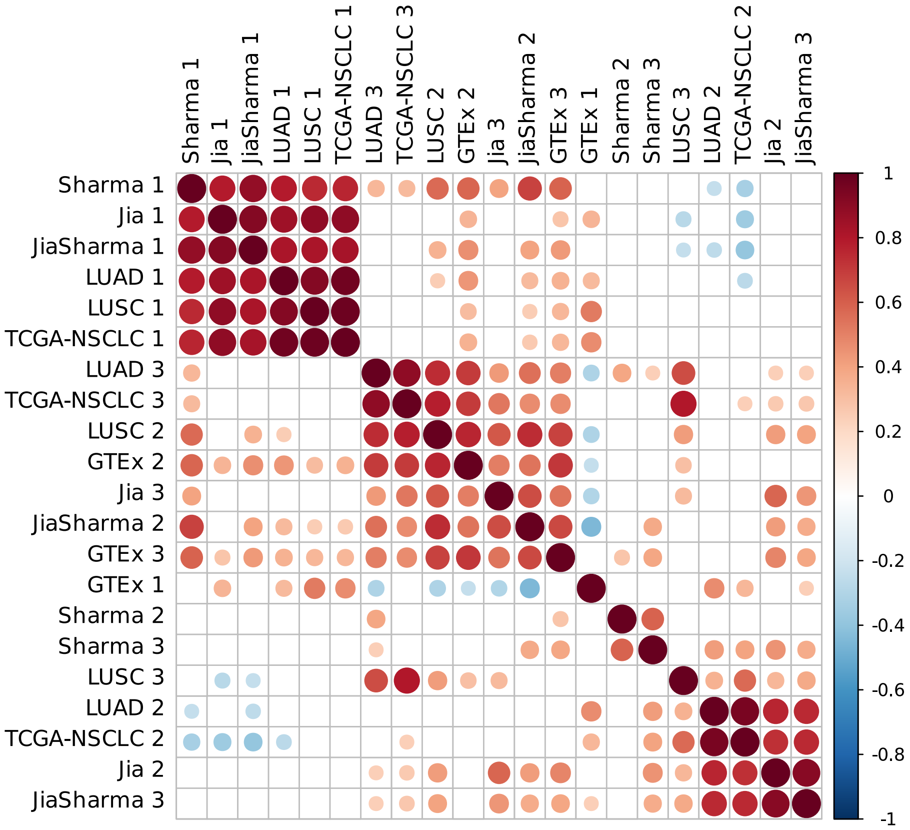


**Figure S2. Correlation heatmap of F1-F3 factors for the different lung-cancer and healthy-lung datasets.** Dot sizes correspond to the absolute correlation coefficient. Only dots where the FDR of the correlation is < 0.01 are shown.

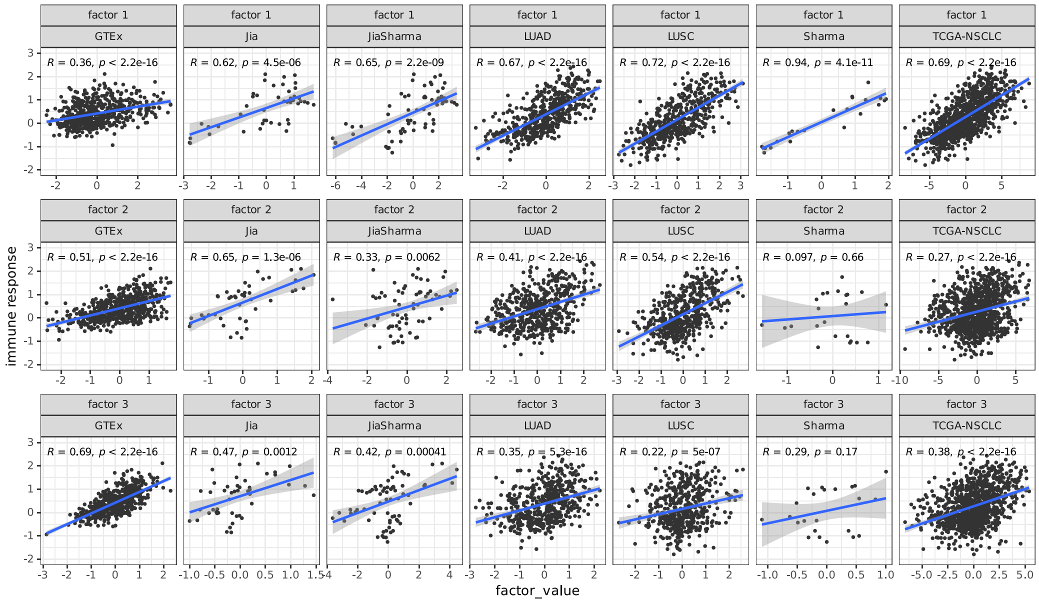


**Figure S3. Correlation scatterplots of MOFA factors with the predicted immune response.** Correlation of F1-F3 factors with the ensemble immune response score, derived from state-of-the-art signatures, for the datasets GTEx, Jia, JiaSharma, LUAD, LUSC, Sharma, and TCGA-NSCLC.

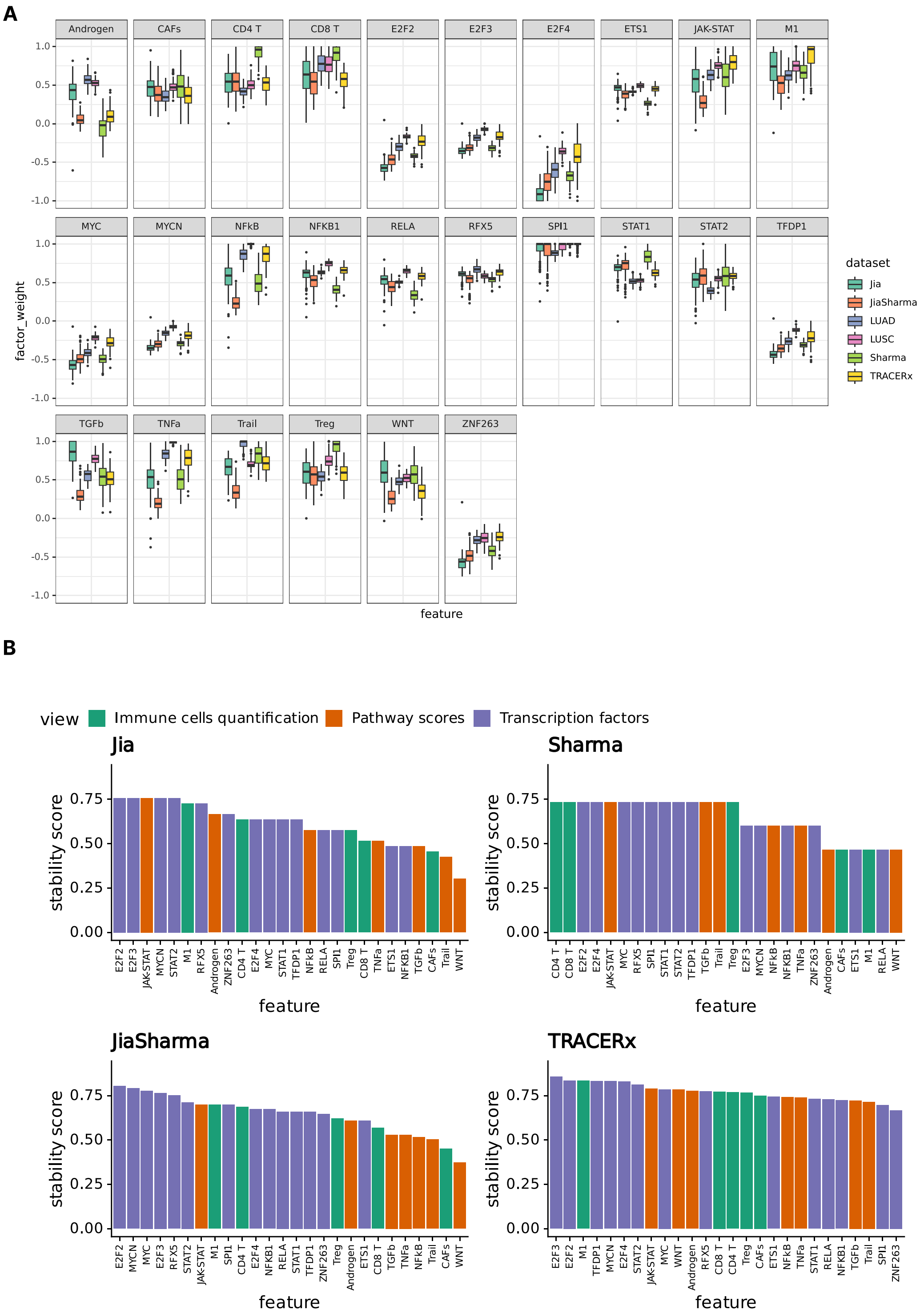


**Figure S4. Feature stability across bootstrap models and tumor biopsies. (A)** MOFA factor weight for top features (as highlighted in Figure 1F) across 100 bootstrap runs. The central line denotes the median. The boxes extend to the 25 and 75 percentiles. The whiskers extend to the most extreme data point within 1.5 of the interquartile range (IQR). **(B)** Rank-based stability score for the Jia, Sharma, JiaSharma and TRACERx datasets. A high feature stability score indicates a low intra-patient heterogeneity.

A

B

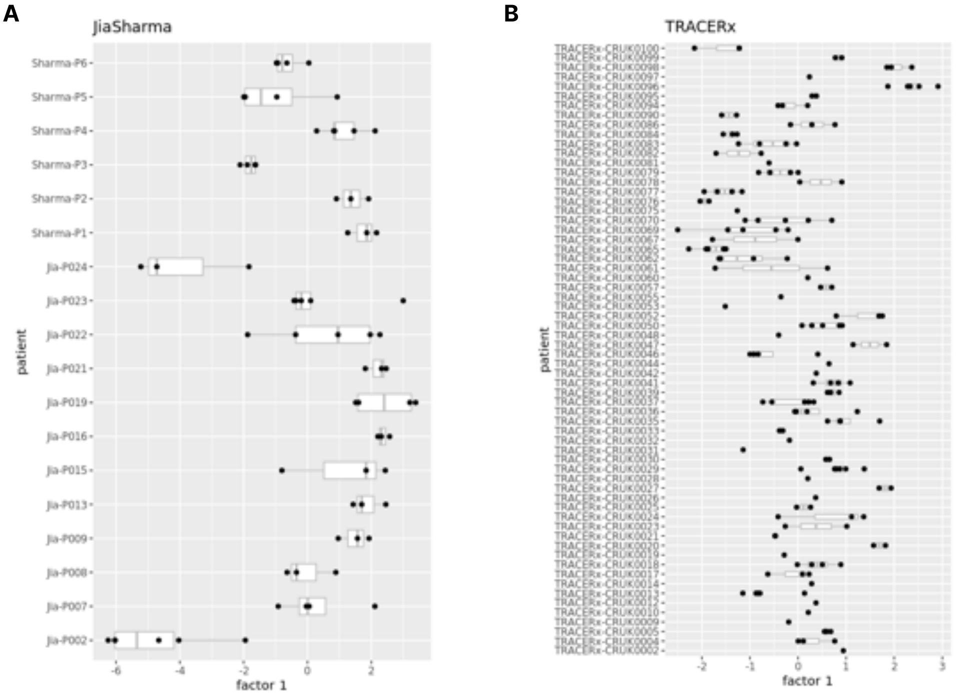


**Figure S5. Intra-tumoral heterogeneity in multi-biopsy data.** For each patient in the JiaSharma and TRACERx datasets, multiple biopsies were taken from different tumor locations. The figure shows the distribution of F1 weights across multiple biopsies for each patient for JiaSharma **(A)** and TRACERx **(B)**. Of the boxplots, the central line denotes the median. The boxes extend to the 25 and 75 percentiles. The whiskers extend to the most extreme data point within 1.5 of the interquartile range (IQR). The black bar denotes the mean of all values.

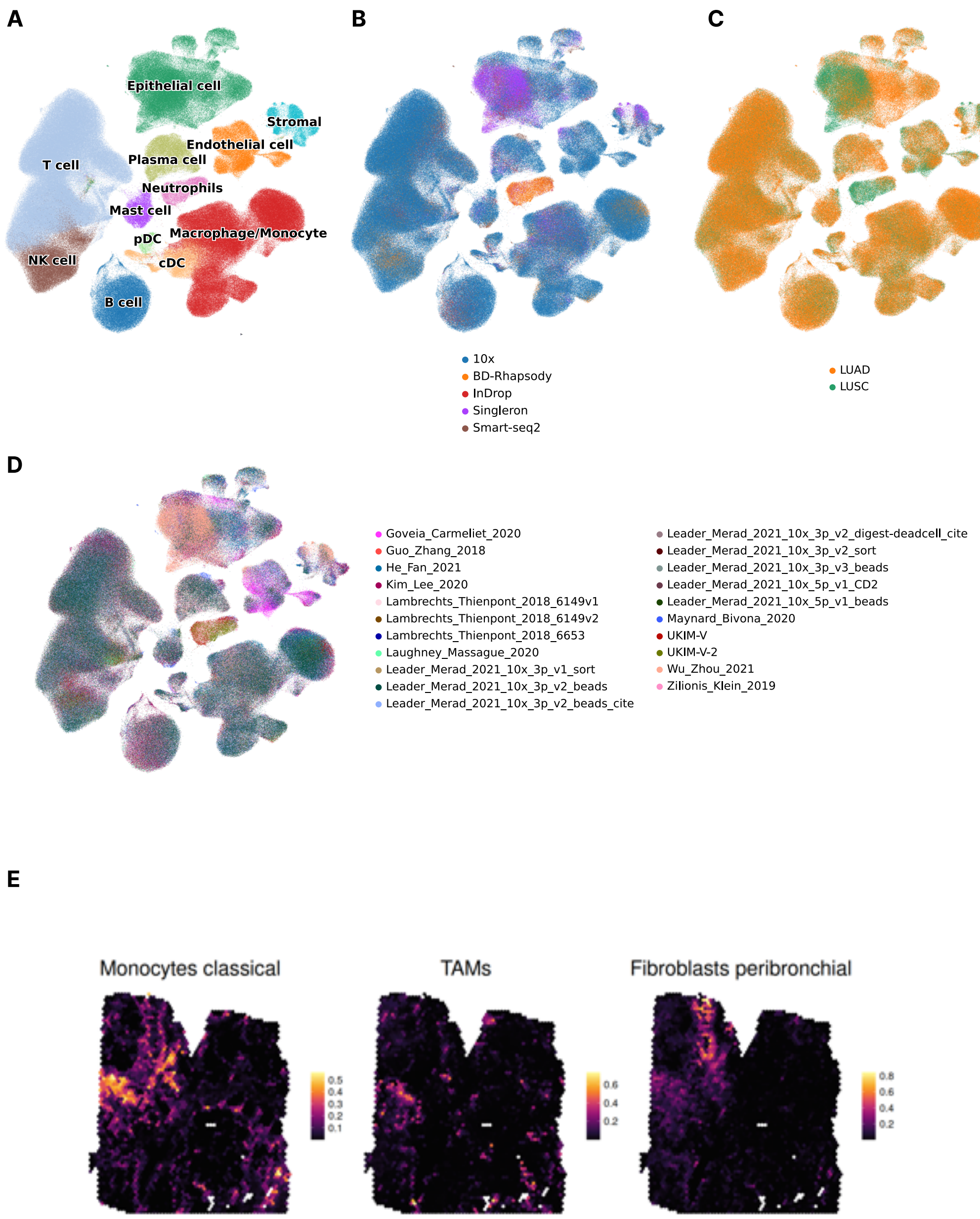


**Figure S6. Covariates of the NSCLC single-cell atlas. (A-D)** UMAP plot of the NSCLC single-cell atlas, colored by **(A)** coarse cell-type labels, **(B)** sequencing platform, **(C)** histological subtype, and **(D)** dataset of origin. **(E)** Exemplary spatial transcriptomics analysis of a lung cancer slide profiled with the 10x Visium technology (same slide as in Figure 2c). The three panels show the estimated cell-type fractions per spot for classical monocytes, tumor-associated macrophages (TAMs), and peribronchial fibroblasts, respectively.

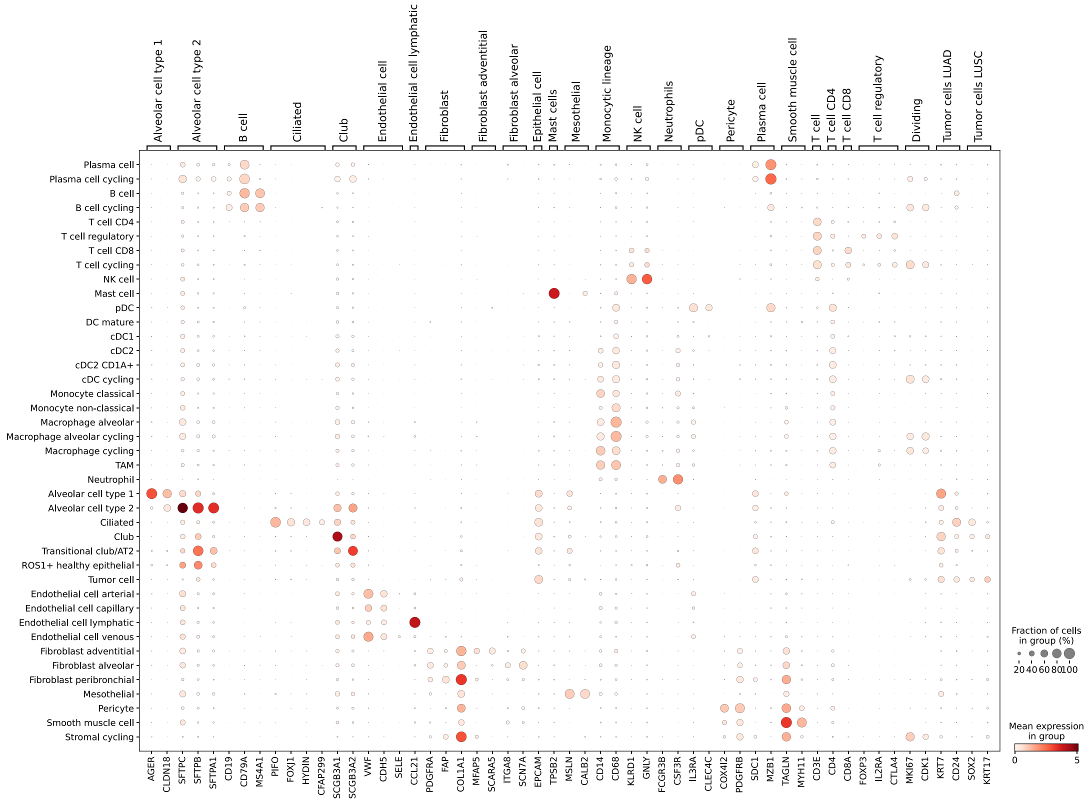


**Figure S7. Cell-type markers.** Dotplot showing the expression of cell-type specific marker genes across the cell-type clusters of the full single-cell dataset.

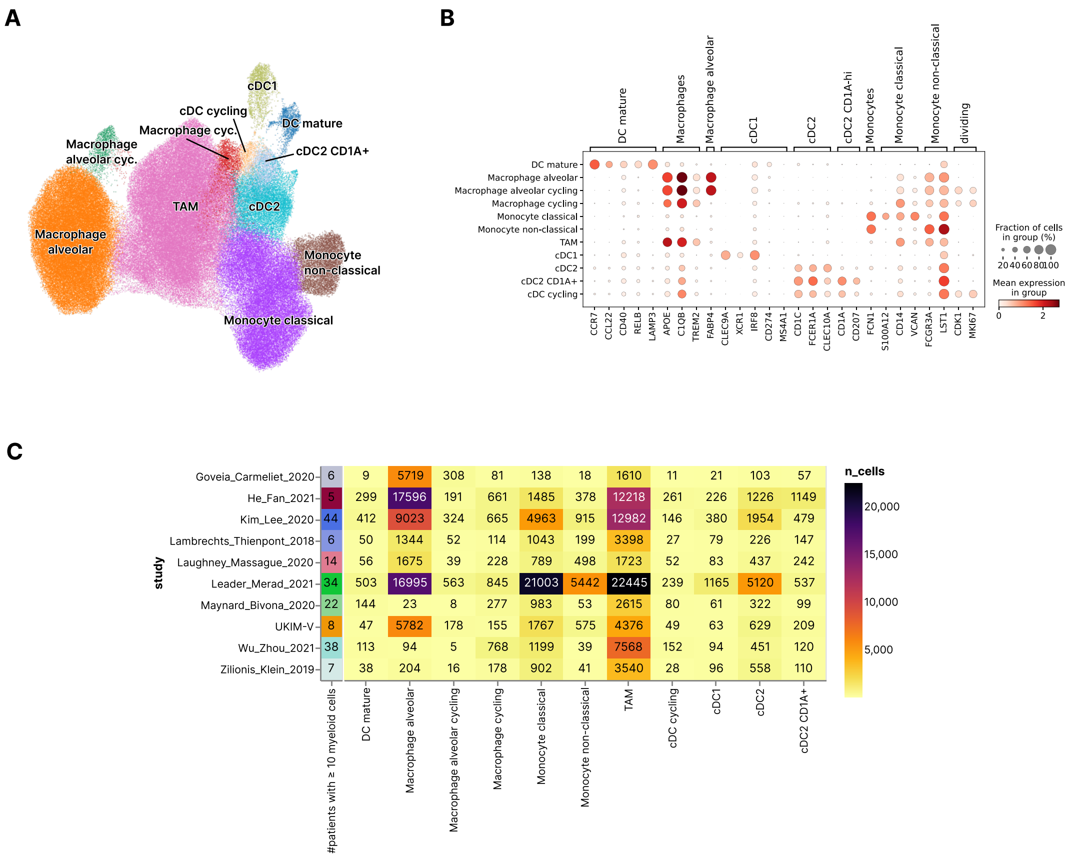


**Figure S8. Analysis of the myeloid subcluster. (A)** UMAP of the myeloid cluster, colored by cell-type. **(B)** Dotplot of marker genes characteristic for myeloid subsets. **(C)** Number of cells per myeloid cell-type and dataset. The first column of the matrix indicates the number of patients per dataset that have at least ten myeloid cells.

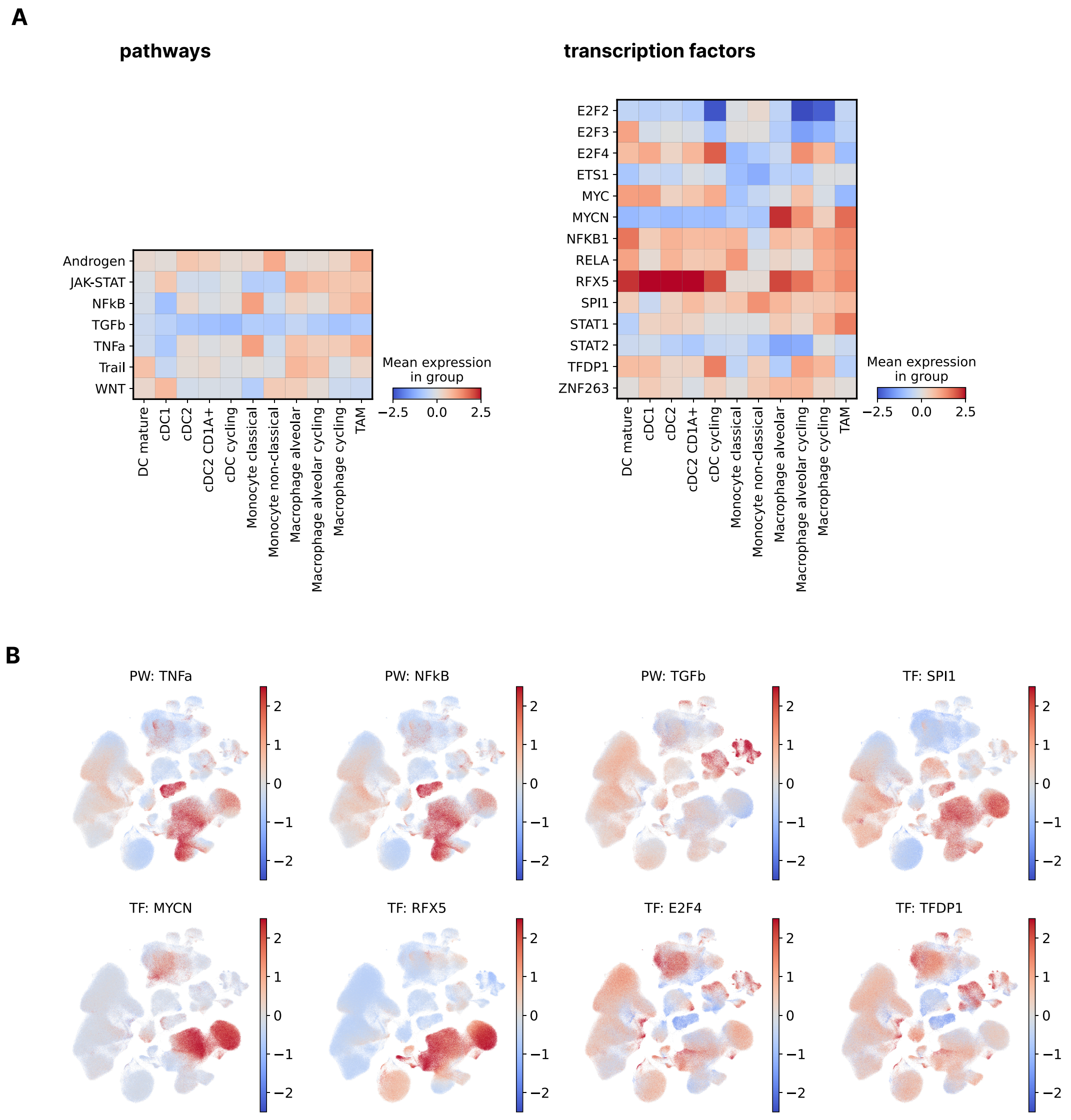


**Figure S9. Pathway and transcription factor activity in myeloid cells. (A)** Heatmap of average activity scores of top pathways and transcription factors (as highlighted in Figure 1F) per myeloid cell-type as determined using Progeny and Dorothea, respectively. Values are z-scores of activity scores computed for each row. **(B)** UMAP plots colored by the activity scores for selected pathways (PW) and transcription factors (TF).

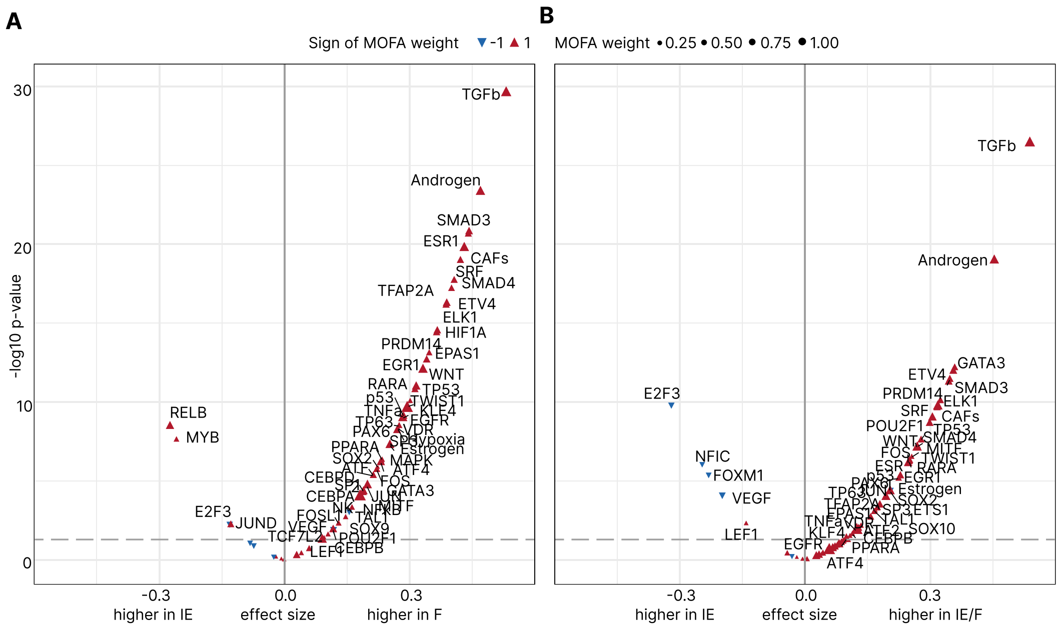


**Figure S10. Comparison of features associated to immune-exclusion in patients from immune-enriched and fibrotic subtypes. (A)** Immune-enriched (IE) vs immune-enriched, fibrotic (IE/F) and **(B)** IE vs fibrotic (F) patients (Wilcoxon rank-sum test). Each dot represents a iHet feature, where the size and color represents the corresponding weight and sign determined by MOFA, respectively.

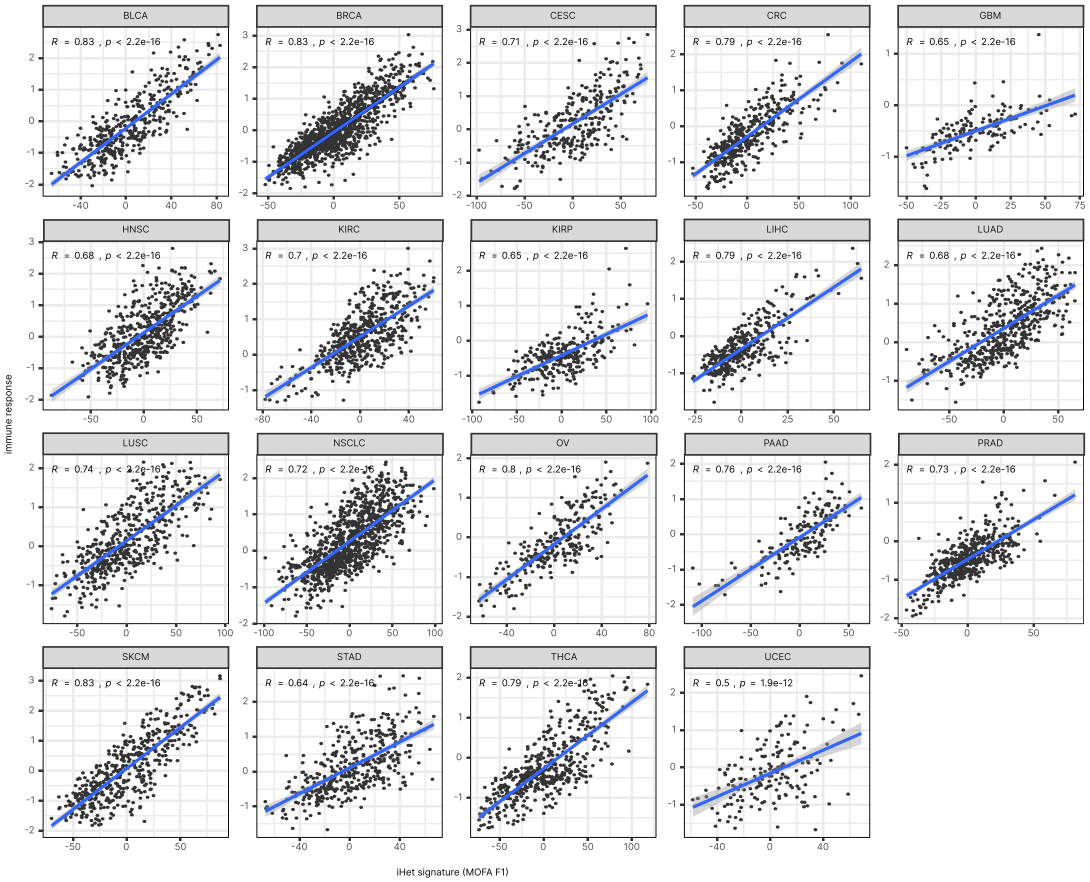


**Figure S11. Correlation of iHet with immune response.** Correlation scatterplots of iHet score (x-axis) with the ensemble immune response score (IR; y-axis) for each TCGA cancer type. *R* represents Pearson correlation, *p* the associated, two-tailed p-value.

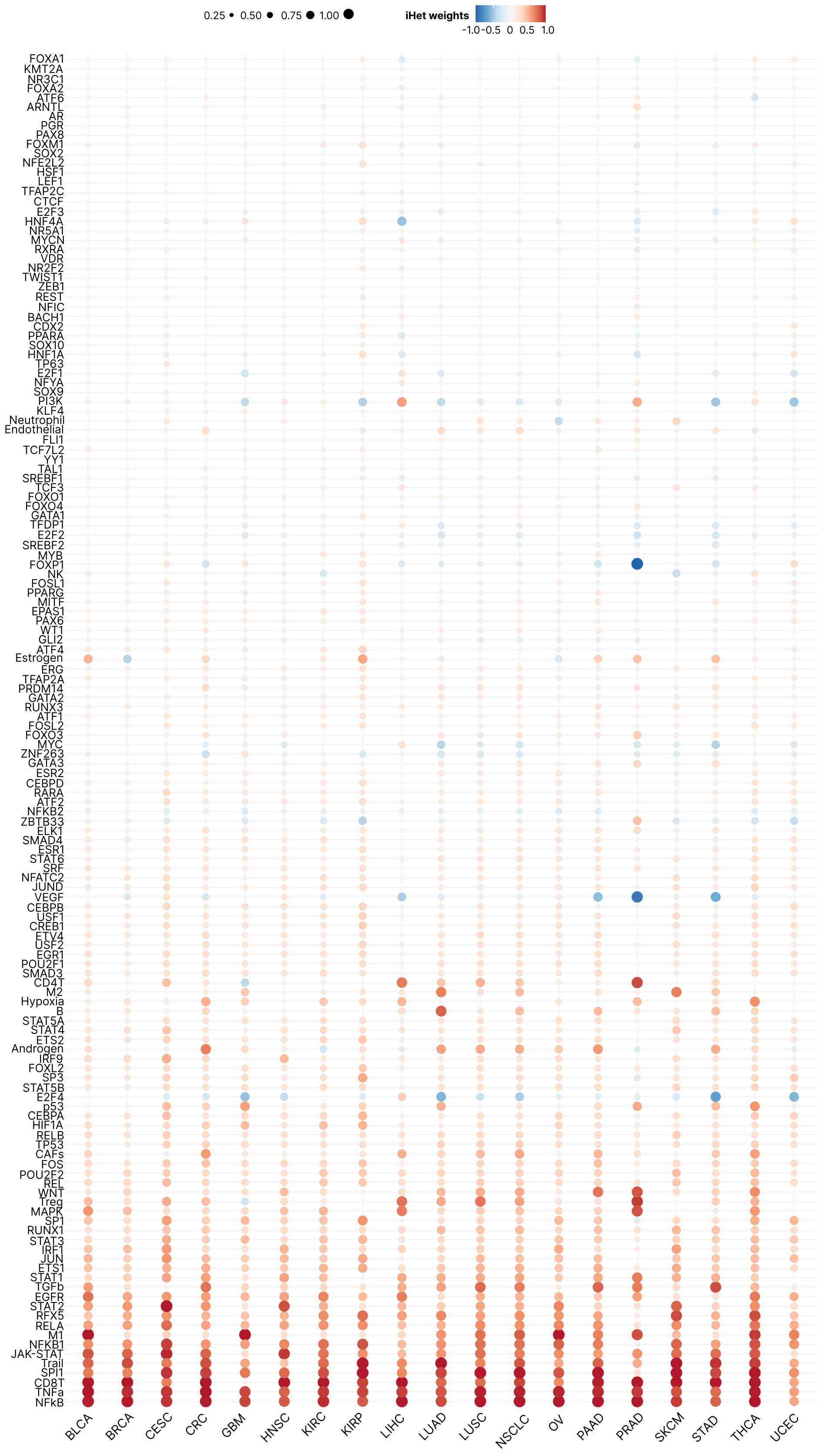


**Figure S12. iHet-associated feature weights.** Heatmap showing cancer-type-specific iHet-associated feature weights. Shown are the median values computed across 100 bootstrap models. Rows (features) were sorted according to their absolute mean value across cancer types. Weights were scaled into the [-1,1] range in each bootstrap model.

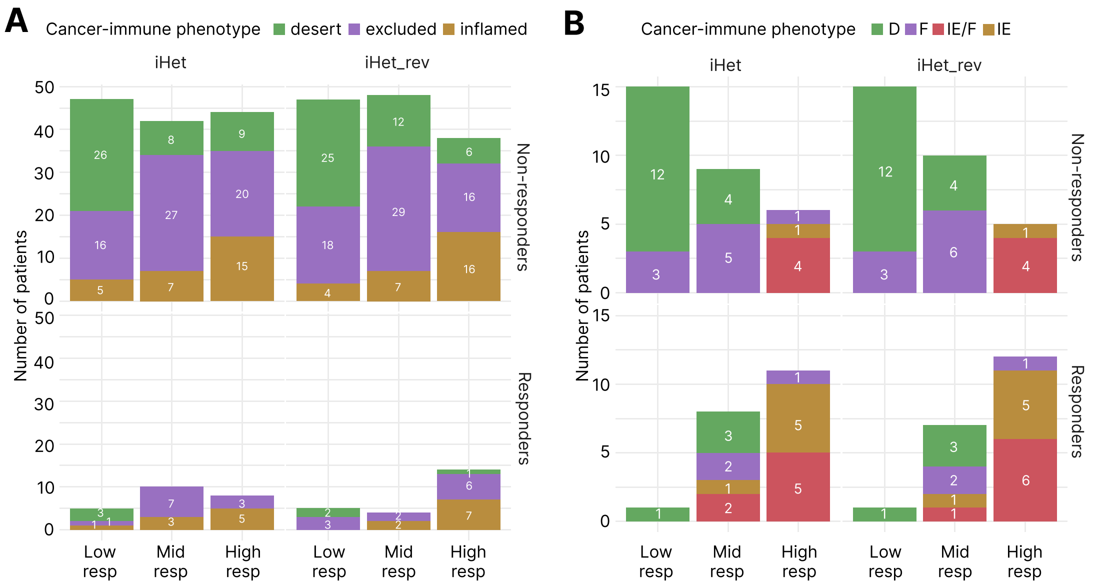


**Figure S13. iHet and iHet_rev classification of patients and association with cancer-immune phenotypes.** Classification of non-responding (left) and responding (right) patients in three tertiles defined using iHet and iHet_rev scores for **(A)** the Mariathasan bladder cancer cohort, and **(B)** the combined Gide and Auslander melanoma cohort. Patients' are coloured according to their defined cancer-immune phenotype[1,2].

**A**

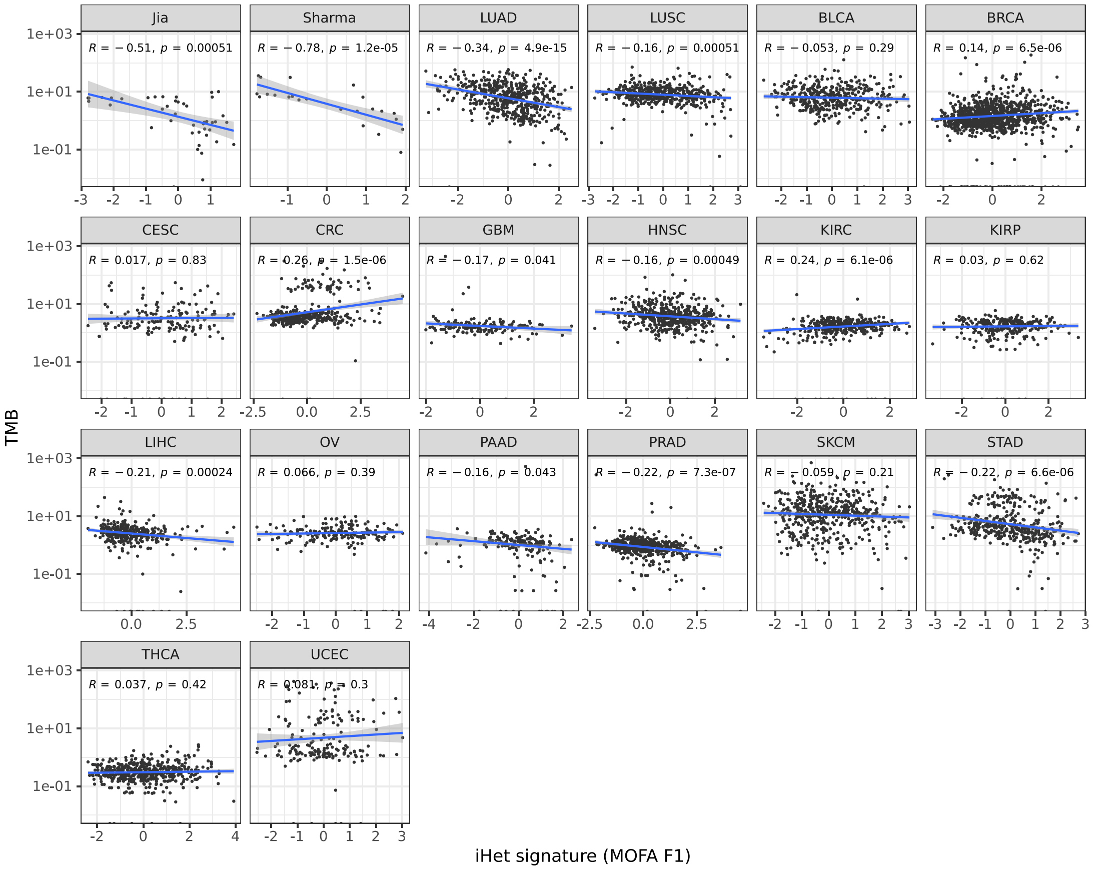


**B**

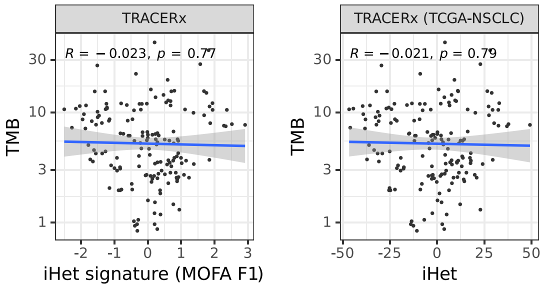


**Figure S14**. **Correlation of iHet with TMB. (A)** Correlation of iHet signature (weights of MOFA F1), on the x-axis, with tumor mutational burden (TMB), on the y-axis, for Jia, Sharma, and all TCGA cohorts. **(B)** Correlation of the iHet signature (left panel) and iHet score (derived from the TCGA-NSCLC signature) with TMB for the TRACERx dataset. *R* represents Pearson correlation, *p* the associated, two-tailed p-value.
